# NEIL1 and NEIL2 DNA glycosylases regulate anxiety and learning in a cooperative manner

**DOI:** 10.1101/2021.02.08.430208

**Authors:** Gunn A. Hildrestrand, Veslemøy Rolseth, Nicolas Kunath, Rajikala Suganthan, Vidar Jensen, Anna M. Bugaj, Marion S. Fernandez-Berrocal, Sunniva Bøe Sikko, Susanne Vetlesen, Anna Kuśnierczyk, Ann-Karin Olsen, Kristine Bjerve Gützkow, Alexander D. Rowe, Wei Wang, Olve Moldestad, Monica Dahl Syrstad, Geir Slupphaug, Lars Eide, Arne Klungland, Pål Sætrom, Luisa Luna, Jing Ye, Katja Scheffler, Magnar Bjørås

## Abstract

Oxidative DNA damage in the brain has been implicated in neurodegeneration and cognitive decline. DNA glycosylases initiate base excision repair (BER), the main pathway for oxidative DNA base lesion repair. NEIL1 and NEIL3 DNA glycosylases alter cognition in mice, the role of NEIL2 remains unclear. Here, we investigate the impact of NEIL2 and its potential overlap with NEIL1 on behavior in single and double knock-out mouse models. *Neil1*^*-/-*^*Neil2*^*-/-*^ mice displayed hyperactivity, reduced anxiety and improved learning. Hippocampal oxidative DNA base lesion levels were comparable between genotypes, no mutator phenotype was found. Impaired canonical repair was thus not the cause of altered behavior. Electrophysiology indicated reduced stratum oriens afferents in the hippocampal CA1 region in *Neil1*^*-/-*^*Neil2*^*-/-*^. Within CA1, NEIL1 and NEIL2 jointly regulated transcription in genes relevant for synaptic function. Thus, we postulate a cooperative function of NEIL1 and NEIL2 in genome regulation beyond canonical BER modulating memory formation and anxiety.

## Introduction

Cells in tissues and organs are continuously subjected to oxidative stress originating both from exogenous and endogenous sources such as reactive oxygen species (ROS), ionizing radiation, UV radiation, and chemicals, amongst others ^1^ The brain is especially susceptible to oxidative stress due to a high metabolic rate, low levels of antioxidant enzymes, and high levels of iron ^2–4^ Thus, repair of oxidative damage in the genome of post-mitotic neurons is supposed to be critical for proper brain function ^5–7^ The hippocampus is a brain area critical for learning and memory formation and is also involved in anxiety ^8–12^ Increasing evidence shows that oxidative stress and defective DNA repair affects the hippocampus and leads to cognitive impairment ^13–16^ Oxidative stress has also been implicated in depression and anxiety ^14,16–18^ In mammalian cells, oxidative DNA damage is predominantly repaired via the base excision repair (BER) pathway reviewed in ^19^ and enzymes in this pathway have been shown to be important for protection against neuronal cell death following induced ischemic brain damage ^20–23^ BER is initiated by DNA glycosylases, which recognize and remove small base lesions (reviewed in ^24,25^). To date, eleven mammalian DNA glycosylases have been identified. NEIL1 and NEIL2 are two of five DNA glycosylases specific for oxidative base lesions and the substrate specificities for these DNA glycosylases are partially overlapping. NEIL1 has broad substrate specificity and removes both pyrimidine- and purine-derived lesions such as 4,6-diamino-5-formamidopyrimidine (FapyA), 2,6-diamino-4-hydroxy-5-formamidopyrimidine (FapyG), guanidionhydantoin (Gh), spiroiminodihydantoin (Sp) and thymine glycol (Tg) from DNA. NEIL2 primarily removes oxidation products of cytosine such as 5-hydroxy-cytosine (5-ohC) and 5-hydroxy-uracil (5-ohU), also excised by NEIL1 ^19,26–28^. *NEIL1* and *NEIL2* mRNA is homogeneously distributed and ubiquitously expressed in human and murine brain, indicating a role of NEIL1 and NEIL2 in DNA maintenance in most areas of the brain ^29^. Previous studies of mice lacking NEIL1 revealed a mild metabolic phenotype, impaired memory retention and defects in olfactory function, as well as increased sensitivity to ischemic brain injury ^23,30,31^. No overt phenotype has been reported for NEIL2-deficient mice, but they were shown to accumulate oxidative damage in transcribed regions of the genome with age ^32^. Recently, we reported no accumulation of DNA damage or mutations and no predisposition to cancer development in mice lacking both NEIL1 and NEIL2 ^33^. In the present study, we used mice deficient in NEIL1 and/or NEIL2 DNA glycosylases to elucidate the roles of these enzymes in behavior and cognition (e.g. activity, anxiety learning and memory) and to study their impact on genome stability, gene expression and electrophysiological features in the hippocampus.

## Results

### Hyperactivity, reduced anxiety and improved learning in *Neil1*^*-/-*^*Neil2*^*-/-*^mice

The functional consequences of inactivating the *Neil1* and *Neil2* genes were investigated by behavioral studies. General activity levels and anxiety were examined in an open field maze (OFM) and an elevated zero maze (EZM), and hippocampal functions such as spatial learning and memory were studied by using the Morris water maze (MWM) (Figure 1A). The *Neil1*^*-/-*^ *Neil2*^*-/-*^ mice displayed hyperactive behavior both in the OFM and the EZM by being more active (Figure 1B and C) and covering an increased distance when exploring the two mazes (Figure 1D and E), compared to single KO and WT mice. In both mazes, they entered the center and open area zones more frequently than the other mice (Figure 1F and G), and in the EZM they also spent more time in the open area zones, which is indicative of reduced anxiety (Figure 1I). All these characteristics were also observed in the single KO mice, however, to a lesser extent. In the MWM, mice of all genotypes swam well and learned the position of the hidden escape platform, as indicated by reduced latencies to escape the water during training, days 1 to 4 (Figure 1J). Surprisingly, the *Neil1*^*-/-*^*Neil2*^*-/-*^ mice displayed increased learning compared to WT mice. The *Neil2*^*-/-*^ mice also showed a tendency to increased learning; nonetheless, this was not statistically significant (Figure 1J). During the first retention trial (day 5), all genotypes showed the same occupancy at the target quadrant (Figure 1K), suggesting no substantial differences in spatial memory in any of the genotypes. During the second retention trial (day 12), the *Neil1*^*-/-*^*Neil2*^*-/-*^ mice became less decisive, shown as decreased preference for the target zone (Figure 1K) and a tendency to search further away from the platform (Figure 1L). This could be indicative of impaired long-term memory. In line with a previous report ^30^, the *Neil1*^*-/-*^ mice weighed significantly more than the WT mice (Figure S1). This did not seem to affect their activity level (Figures 1B-E). No weight differences were observed for the *Neil2*^*-/-*^ and *Neil1*^*-/-*^*Neil2*^*-/-*^ mice relative to WT mice. Overall, the behavioral tests revealed hyperactivity, reduced anxiety-like behavior, and improved learning in the *Neil1*^*-/-*^*Neil2*^*-/-*^ mice. The single knockouts were only moderately affected.

**Figure 1.**
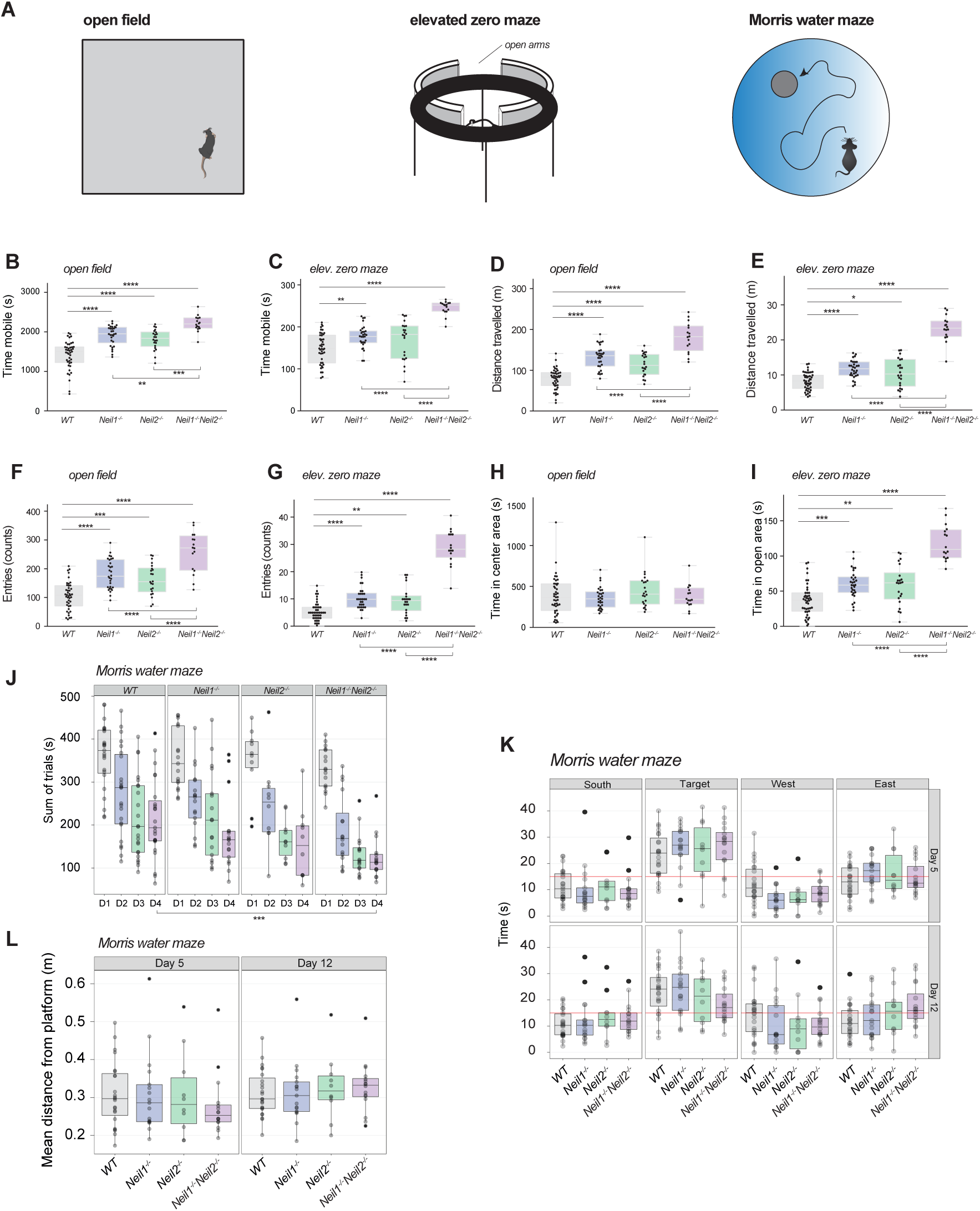
Increased activity, reduced anxiety and enhanced learning in *Neil1*^*-/-*^*Neil2*^*-/-*^mice. In the open field test (A), mice were allowed to explore freely for 45 minutes in an arena measuring L40 cm x W40 cm x H35 cm. An area of L20 cm x W20 cm was defined as the center area zone. (B) Time mobile, (D) distance travelled, (F) entries to the center area zone, and (H) time in the center area zone. In the elevated zero maze (A), the mice were allowed 5 minutes for exploration on a 5-cm wide circular runway with alternating open and closed areas. (C) Time mobile, (E) distance travelled, (G) entries to the open area zones, and (I) time in the open area zones. (B-I) Data are shown in full, with overlaid boxplots representing the medians and the interquartile ranges (IQR). Whiskers indicate min/max values. n = 43 WT, 30 *Neil1*^*-/-*^, 22 *Neil2*^*-/-*^ and 16 *Neil1*^*-/-*^*Neil2*^*-/-*^ mice. * p < 0.05, ** p < 0.01, *** p < 0.001, **** p < 0.0001 by one-way ANOVA/Tukey. In the Morris water maze (A), mice were trained to locate an escape platform hidden below the water surface (days 1-4) before memory was tested (days 5 and 12). (J) Latency to locate the platform and escape the water during learning trials, days 1 to 4. The data are shown as the total time spent in the tank for each mouse during 8 trials. *** p = 0.00019 for *Neil1*^*-/-*^*Neil2*^*-/-*^ vs. WT by non-parametric pair wise Wilcoxon rank-sum test, Holm-adjusted. (K) Time spent in the four quadrants of the tank (the red line indicates average level (15 sec) expected by random behavior) and (L) mean distance from the platform zone during retention trials (probe tests), days 5 and 12. (J-L) Data are shown in full, with overlaid boxplots representing the medians and the interquartile ranges (IQR). Whiskers extend to a Tukey fence set at 1.5xIQR. n = 22 WT, 17 *Neil1*^*-/-*^, 10 *Neil2*^*-/-*^ and 16 *Neil1*^*-/-*^*Neil2*^*-/-*^ mice.

### No change in steady-state levels of oxidative DNA base lesions in *Neil1*^*-/-*^*Neil2*^*-/-*^hippocampus but decreased levels of single-strand breaks

As hippocampus is one of the critical brain areas involved in anxiety as well as learning and memory, we assessed the effect of NEIL1 and/or NEIL2 deficiencies on hippocampal DNA integrity. We applied three different methods. First, the bulk level of the oxidative DNA base lesion 5-ohC, a substrate for both NEIL1 and NEIL2, in hippocampal genomic DNA was measured by mass spectrometry. The results showed no significant differences in global 5-ohC levels between the four genotypes (Figure 2B). Second, the alkaline comet assay was used to study DNA damage, including strand breaks, at the single-cell level. The mutants did not accumulate more DNA damage than WT mice; on the contrary, the hippocampi of *Neil2*^*-/-*^ and *Neil1*^*-/-*^*Neil2*^*-/-*^ mice displayed reduced levels of strand breaks genome-wide as compared to WT mice (Figure 2C). Fpg treatment, unravelling unrepaired base lesions, did not lead to increased comet tail lengths in any of the genotypes, supporting the MS data showing unchanged global levels of oxidative DNA base lesions in all three mutants compared to WT (Figure 2C). Third, a site-specific restriction enzyme-based qPCR method was applied to measure DNA lesions (i.e. AP-sites, base lesions, strand breaks) or mutations. No significant changes were found, but as for the comet assay, there was a tendency to reduction in lesions/mutations in *Neil2*^*-/-*^ and *Neil1*^*-/-*^*Neil2*^*-/-*^ hippocampus, as compared to WT (Figure S2). RNA sequencing analysis revealed no significant changes in the expression of DNA glycosylases with overlapping substrate specificities, such as *Ogg1, Neil3* and *Nth1* (Figure S4 D-F), indicating that there is no compensatory upregulation of these repair genes in mice lacking NEIL1 and/or NEIL2. In sum, our data suggest that NEIL1 and/or NEIL2 do not contribute to the genome-wide repair of DNA in the hippocampus of adult mice.

**Figure 2.**
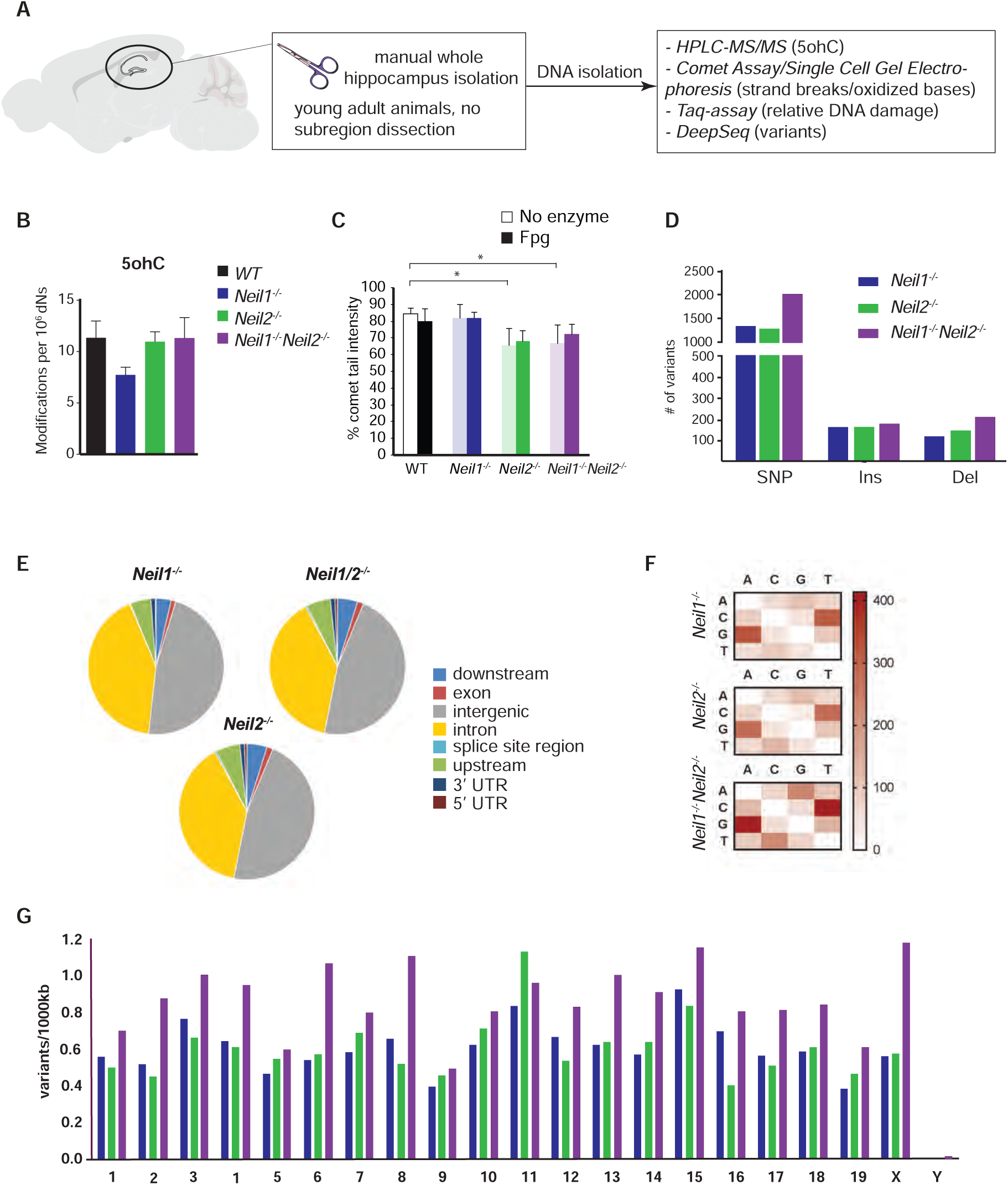
Unchanged steady-state levels of oxidative DNA base lesions and no hypermutator phenotype in *Neil1*^*-/-*^*Neil2*^*-/-*^hippocampus. (A) Hippocampus was isolated from WT and NEIL-deficient mice and DNA damage and mutation levels estimated by various methods. (B) HPLC-MS/MS analysis of 5-ohC in hippocampal, genomic DNA. Data are shown as mean + SEM. n = 5 WT, 5 *Neil1*^*-/-*^, 10 *Neil2*^*-/-*^ and 6 *Neil1*^*-/-*^*Neil2*^*-/-*^ mice. (C) DNA damage levels in hippocampal tissue by alkaline comet assay analysis. Data are shown as mean of gel medians (50 comets x 3 gels scored per mouse) + SEM. n = 4 mice from each genotype. * p = 0.0291 and 0.0495 for *Neil2*^*-/-*^ and *Neil1*^*-/-*^*Neil2*^*-/-*^ vs. WT, respectively, by two-way ANOVA/Sidak. (D-G) DNA samples from four mice of each genotype were pooled and subjected to whole-genome deep sequencing followed by mutation profile analysis. (D) DNA sequence variants, (E) Genomic region distribution of DNA sequence variants, (F) Base changes count of SNPs, and (G) Chromosomal distribution of DNA sequence variants in NEIL-deficient vs. WT hippocampus. SNP, single nucleotide polymorphism; Ins, insertions and Del, deletions.

### No hypermutator phenotype in *Neil1*^*-/-*^*Neil2*^*-/-*^hippocampus

Although the total steady-state levels of mutagenic oxidative DNA lesions were unaltered in hippocampi from adult mice lacking NEIL1 and/or NEIL2, mutations accumulated during development could still explain the phenotype at adulthood. To test this possibility, we applied whole-genome deep sequencing of hippocampal DNA to determine mutation profiles. A DNA sequence variant analysis was performed using the WT hippocampus sample as reference genome. We found a modest increase in DNA sequence variants genome-wide that were evenly distributed across all chromosomes in all the three mutants (Figure 2D and G). Variants were detected in all genomic regions with the majority occurring in non-coding regions, such as intergenic regions and introns (Figure 4E). Analysis of base pair changes in SNPs showed a normal distribution with C:G to T:A transitions being the most frequent, most likely due to deamination of 5mC and C to thymine and uracil, respectively (Figure 4F). These results suggest that lack of NEIL1 and/or NEIL2 does not lead to a genome-wide hypermutator phenotype in hippocampus.

### Reduction in fiber density or number of afferent fibers in stratum oriens of *Neil1*^*-/-*^*Neil2*^*-/-*^hippocampus

To assess potential changes in excitatory synaptic transmission and cell excitability tested by synaptic activation, that could possibly explain the altered behavior in NEIL1/NEIL2-deficient mice, we recorded in either stratum radiatum (SR) or stratum oriens (SO) and simultaneously in stratum pyramidale (SP) in the CA1 region of hippocampal slices from *Neil1*^*-/-*^*Neil2*^*-/-*^ and WT mice. We decided to focus on the hippocampal CA1 subfield due to its prominent role in both spatial information coding and anxiety regulation (Bannerman et al., 2014; Buzsáki and Moser, 2013; Jimenez et al., 2018; O’Keefe and Nadel, 1978).

The stimulation strengths necessary to elicit prevolleys of given amplitudes (0.5, 1.0 and 1.5 mV) tended to be higher, though not statistically significant, in SR of *Neil1*^*-/-*^*Neil2*^*-/-*^ mice compared to WT mice (Figure 3A, i). In SO, on the other hand, significantly higher stimulation strengths were needed in *Neil1*^*-/-*^*Neil2*^*-/-*^ mice compared to WT mice (Figure 3A, vi). Measuring the field excitatory postsynaptic potential (fEPSP) as a function of the same prevolley amplitudes showed that *Neil1*^*-/-*^*Neil2*^*-/-*^ animals evoked fEPSPs similar to those obtained in WT mice, in both SR (Figure 3A, ii and v) and SO (Figure 3A, vii and x). Furthermore, postsynaptic excitability, measured as fEPSPs necessary for generating a population spike, was not significantly changed in *Neil1*^*-/-*^*Neil2*^*-/-*^ mice compared to WT mice in SR (Fig 3A, iii and v) or SO (Figure 3A, viii and x).

**Figure 3.**
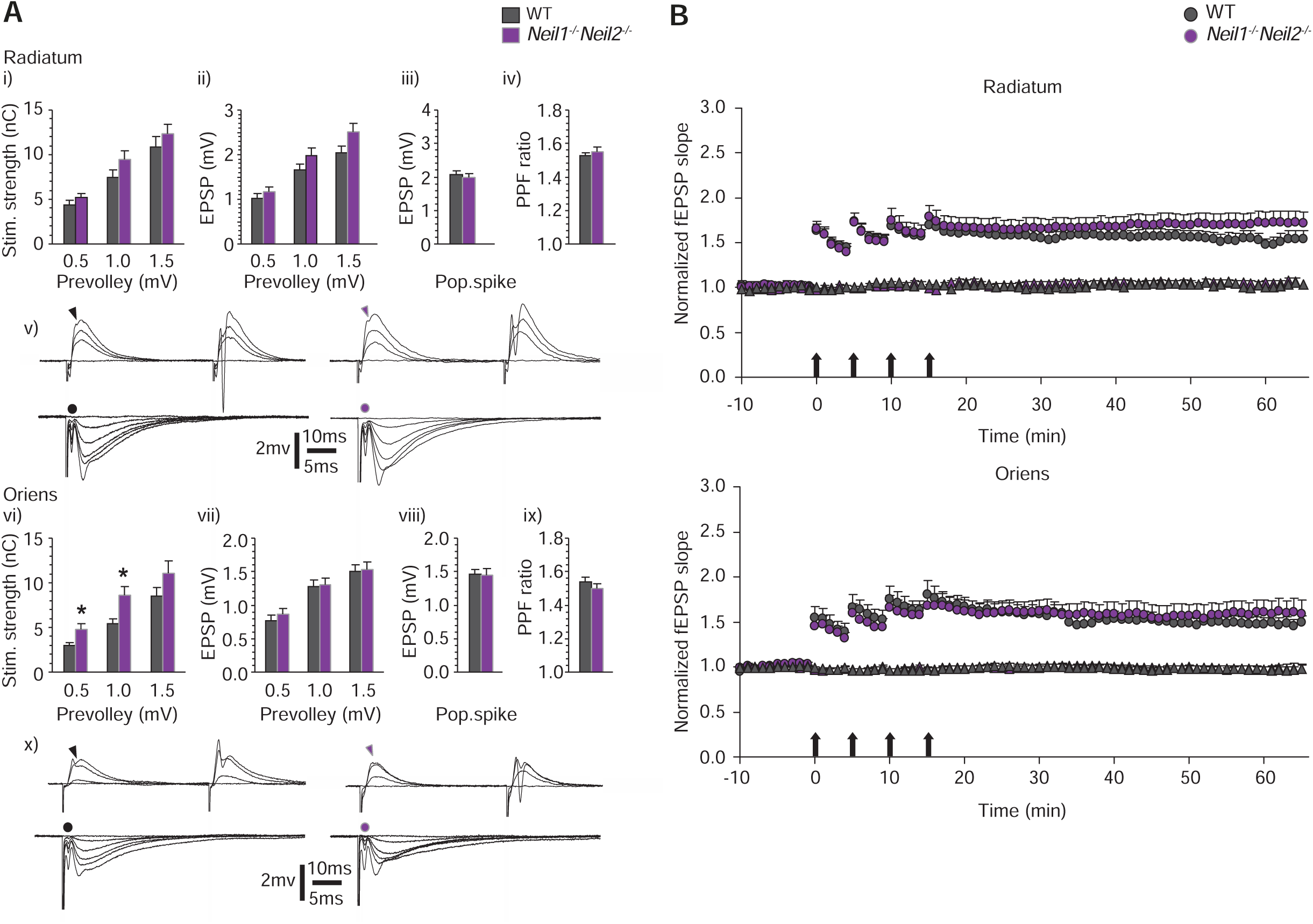
Reduced number of afferent fibers in stratum oriens of *Neil1*^*-/-*^*Neil2*^*-/-*^hippocampus. (A) Synaptic transmission, excitability and paired-pulse facilitation (PPF) in stratum Radiatum (SR) (i – v) and stratum oriens (SO) (vi – x) of *Neil1*^*-/-*^*Neil2*^*-/-*^ and WT mice. (i) and (vi) Stimulation strengths necessary to elicit prevolleys of given amplitudes (0.5, 1.0 and 1.5 mV). (ii) and (vii) fEPSP amplitudes as a function of the same three prevolley amplitudes. (iii) and (viii) The fEPSP amplitudes necessary to elicit a just detectable population spike. (iv) and (ix) PPF ratio from the two genotypes at an interstimulus interval of 50 ms. (v) and (x) Top row; recordings from stratum pyramidale elicited by paired-pulse stimulation (50 ms interstimulus interval) in control (left) and *Neil1*^*-/-*^*Neil2*^*-/-*^ (right) mice. Arrowheads indicate the population spike thresholds. Bottom row; each trace is the mean of five consecutive synaptic responses elicited by different stimulation strengths in slices from control (left) and *Neil1*^*-/-*^*Neil2*^*-/-*^ (right) mice. The prevolleys preceding the fEPSPs are indicated by circles. Data are shown as mean + SEM. (i) and (ii) n = 32, 32, 28 WT and 32, 32, 29 *Neil1*^*-/-*^*Neil2*^*-/-*^ for 0.5, 1.0, and 1.5 mV, respectively; (iii) and (iv) n = 32 for both genotypes; (vi) and (vii) n = 28, 28, 25 WT and 32, 31, 26 *Neil1*^*-/-*^*Neil2*^*-/-*^ for 0.5, 1.0, and 1.5 mV, respectively; (viii) and (ix) n = 28 WT and 32 *Neil1*^*-/-*^*Neil2*^*-/-*^. * p = 0.019 and 0.022 for prevolleys of 0.5 and 1.0 mV, respectively, by linear mixed model analysis. (B) Normalized and pooled fEPSP slopes evoked in hippocampal slices from WT and *Neil1*^*-/-*^*Neil2*^*-/-*^ mice in SR and SO. The tetanized pathways are shown as circles and the untetanized control pathways are shown as triangles. Arrows indicate time points of tetanic stimulation. Data are shown as mean + SEM. SR, n = 16 for both genotypes; SO, n = 17 WT and 15 *Neil1*^*-/-*^*Neil2*^*-/-*^. (p = 0.25 and 0.45 for *Neil1*^*-/-*^*Neil2*^*-/-*^ vs. WT in SR and SO, respectively, by linear mixed model analysis).

In sum, the results do not support any major differences between the mutant and WT mice in excitatory synaptic transmission (ii and vii) or postsynaptic excitability (iii, viii) in either of the two strata examined. However, in SO (vi), but not in SR (i), altered axonal activation suggests a reduction in fiber density or number of afferent fibers in in *Neil1*^*-/-*^*Neil2*^*-/-*^ mice compared to WT.

### Similar short-term plasticity in the CA1 region in *Neil1*^*-/-*^*Neil2*^*-/-*^and WT mice

To further characterize excitatory synaptic transmission in the hippocampal CA1 region, we measured paired-pulse facilitation (PPF) ^36^, a short-lasting form of synaptic plasticity primarily attributed to changes in presynaptic Ca^2+^ homeostasis ^37^. A comparison of PPF did not reveal any differences between the two genotypes in SR (Figure 3A, iv) or in SO (Figure 3A, ix).

A 20 Hz stimulus train, which activates the afferent fibres in the CA3 to CA1 pathway, induced an early frequency facilitation, which may depend on the size of the readily releasable vesicle pool ^38,39^, and a delayed response enhancement (DRE) appearing 4 – 6 seconds after train initiation and is presumably caused by recruitment of vesicles from endocytotic recycling and/or reserve vesicles ^40,41^. In the present study, we observed no differences between the two genotypes in the time needed from the initiation of the stimulation train to reach the peak of the initial frequency facilitation (Figure S3, i), the time to the transition point between the frequency facilitation and the DRE (Figure S3, ii), or the time to the maximal value of DRE (Figure S3, iii).

### Similar amount of LTP at CA3 to CA1 synapses in *Neil1*^*-/-*^*Neil2*^*-/-*^and WT mice

We next analyzed long-term potentiation of synaptic transmission (LTP) at CA3 to CA1 synapses in WT and *Neil1*^*-/-*^*Neil2*^*-/-*^ mice in SR and SO. Tetanic stimulation of the afferent fibers in either of the pathways produced a lasting, homosynaptic potentiation of the fEPSP slope of similar magnitude in *Neil1*^*-/-*^*Neil2*^*-/-*^ and control mice, when measured 40 – 45 min after the tetanizations (Figure 3B). In *Neil1*^*-/-*^*Neil2*^*-/-*^ mice, LTP in both SR and SO were similar in magnitude to the corresponding pathways in WT mice.

### Differential expression of genes regulating synaptic function, plasticity and composition

We recently reported hippocampal transcriptional changes in mice lacking OGG1 and MUTYH DNA glycosylases ^16^. We therefore asked whether NEIL1 and NEIL2 glycosylases could similarly act as transcriptional regulators within the hippocampal CA1 region to modulate synaptic transmission and behavior. We applied whole genome sequencing of RNA isolated from the pyramidal layer of CA1 by laser capture microdissection (Figure 4A), a method which offers supreme tissue specificity ^42^. A moderate number of differentially expressed genes (DEGs) were detected, the largest amount in *Neil2*^*-/-*^ and *Neil1*^*-/-*^*Neil2*^*-/-*^ mice (Figure 4B, Figure S4D-F). Similar numbers of up- and downregulated genes were found in all genotypes (Figure 4C). While there was almost no overlap in DEGs between *Neil1*^*-/-*^ and *Neil2*^*-/-*^, we found most overlapping DEGs between single- and double-knockout mice (Figure 4D). A reactome pathway analysis showed the nuclear receptor signaling pathway (R-MMU-38328) to be significantly overrepresented in *Neil1*^*-/-*^*Neil2*^*-/-*^ CA1. Of note, all three isotypes of the orphan nuclear receptor Nr4a were downregulated in *Neil1*^*-/-*^*Neil2*^*-/-*^ mice, whereas the nuclear receptors Nr1d1 and Nr1d2 were upregulated (Figure 4E). While four of these five nuclear receptors were similarly differentially regulated in *Neil2*^*-/-*^, none of them were altered in *Neil1*^*-/-*^ CA1, pointing to a NEIL2-dependent regulation of nuclear receptors. The top10 upregulated genes in *Neil1*^*-/-*^*Neil2*^*-/-*^ mice largely overlapped with those of *Neil2*^*-/-*^ mice, whereas downregulated genes overlapped mainly with those of *Neil1*^*-/-*^ mice (Figure 4F). Among the up- and downregulated DEGs we identified four genes as immediately relevant to synaptic function according to their QuickGO annotation ^43^ (Figure 4F). While three of them (Npbwr1, Tmem254b and Fxyd2) play a role in a very specific subset of receptor systems and synaptic membrane elements, Npas4 is a well-characterized master regulator of inhibitory synapse development ^44^. The latter was differentially regulated distinctly in *Neil1*^*-/-*^ and *Neil1*^*-/-*^ *Neil2*^*-/-*^ animals, but not in *Neil2*^*-/-*^ animals, indicating a mainly NEIL1-dependent regulation of this gene (Figure 4F). Differences observed in group comparisons were also visible at a single animal level (Figure 4G).

**Figure 4.**
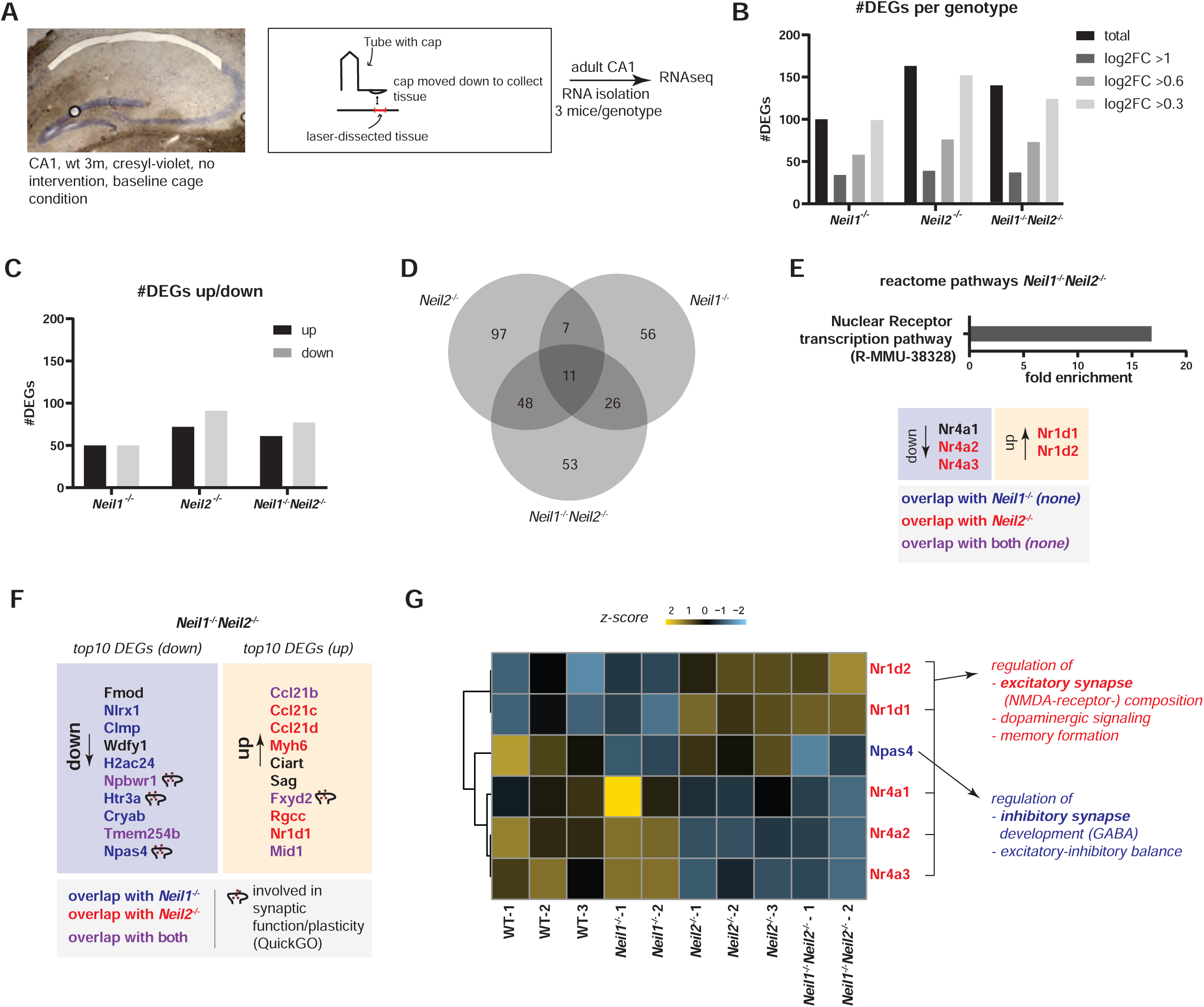
NEIL1 and NEIL2 jointly regulate the CA1 transcriptome. (A) Tissue used for RNAseq was isolated using a laser dissection approach. (B) Amount of DEGs found for each genotype. DEGs above each log2fold cut-off are displayed in different shades of grey. (C) Up-and downregulated genes for each mutant. (D) Overlapping DEGs between single and double knockouts. (E) Enriched reactome pathway, R-MMU-38328 (fold enrichment 19.72, FDR 1.19e-02), and corresponding DEGs in *Neil1*^*-/-*^*Neil2*^*-/-*^ mice. Overlap with *Neil1*^*-/-*^ or *Neil2*^*-/-*^ mice is indicated. (F) top10 up- and downregulated DEGs in *Neil1*^*-/-*^*Neil2*^*-/-*^ mice. (Npas4: -1.12 log2fold). Overlap with *Neil1*^*-/-*^ or *Neil2*^*-/-*^ mice is indicated. (G) Expression of relevant DEGs at a single animal level (z-score table based on FPKMs). Thematic relevance of DEGs in *Neil1*^*-/-*^*Neil2*^*-/-*^ mice is shown to the right.

To further thematically cluster the DEGs found in the different genotypes, we performed a gene ontology biological processes enrichment analysis (PANTHER release 2020-07-28, GO database release 2020-07-16, DEGs log2fold(abs) >0.3, p<0.05). All NEIL-deficient mice showed an enrichment of several GO-terms immediately relevant to central nervous system function (colored GO-terms, Figure S4 A-C), further highlighting the relevance of NEIL DNA glycosylases in CA1 transcription regulation.

### Altered synaptic composition in *Neil2*^*-/-*^and *Neil1*^*-/-*^*Neil2*^*-/-*^mice

Based on the transcriptome results showing differential regulation of factors relevant for synaptic composition, we decided to examine the excitatory and inhibitory transmitter systems by immunohistochemistry within the CA1 subregion of the hippocampal formation. We chose to study the NMDA- and GABA-receptors due to their reciprocal interaction with both Npas4 and Nr4a-isoforms.

Within the tetrameric structure of the NMDA-receptor complex (Figure 5, illustration), regulatory subunits such as NR2A (GRIN2A) and NR2B (GRIN2B) determine the receptor’s electrophysiological properties and are seen as important mediators of synaptic plasticity ^45^. We therefore primarily examined these two subunits of the NMDA receptor. Across stratum pyramidale (SP), we found significantly reduced NR2A-reactivity in *Neil1*^*-/-*^*Neil2*^*-/-*^ mice and *Neil2*^*-/-*^ mice compared to WT (Figure 5A). NR2A reactivity was also significantly lower within stratum oriens (SO) in *Neil1*^*-/-*^*Neil2*^*-/-*^ mice compared to WT. As for the NR2B subunit, reduced reactivity was observed across SP of *Neil2*^*-/-*^ mice only (Figure 5B). A low NR2A/NR2B-ratio has previously been reported to enhance both memory formation ^46^ and LTP ^47^. We observed a significantly reduced NR2A/NR2B ratio exclusively in *Neil1*^*-/-*^*Neil2*^*-/-*^ mice, with the most prominent reduction across SO (SO, Δ2.558; SR, Δ1.685; SP, Δ1.053) (Figure 5C), the region that showed significant electrophysiological differences (Figure 3, vi).

**Figure 5.**
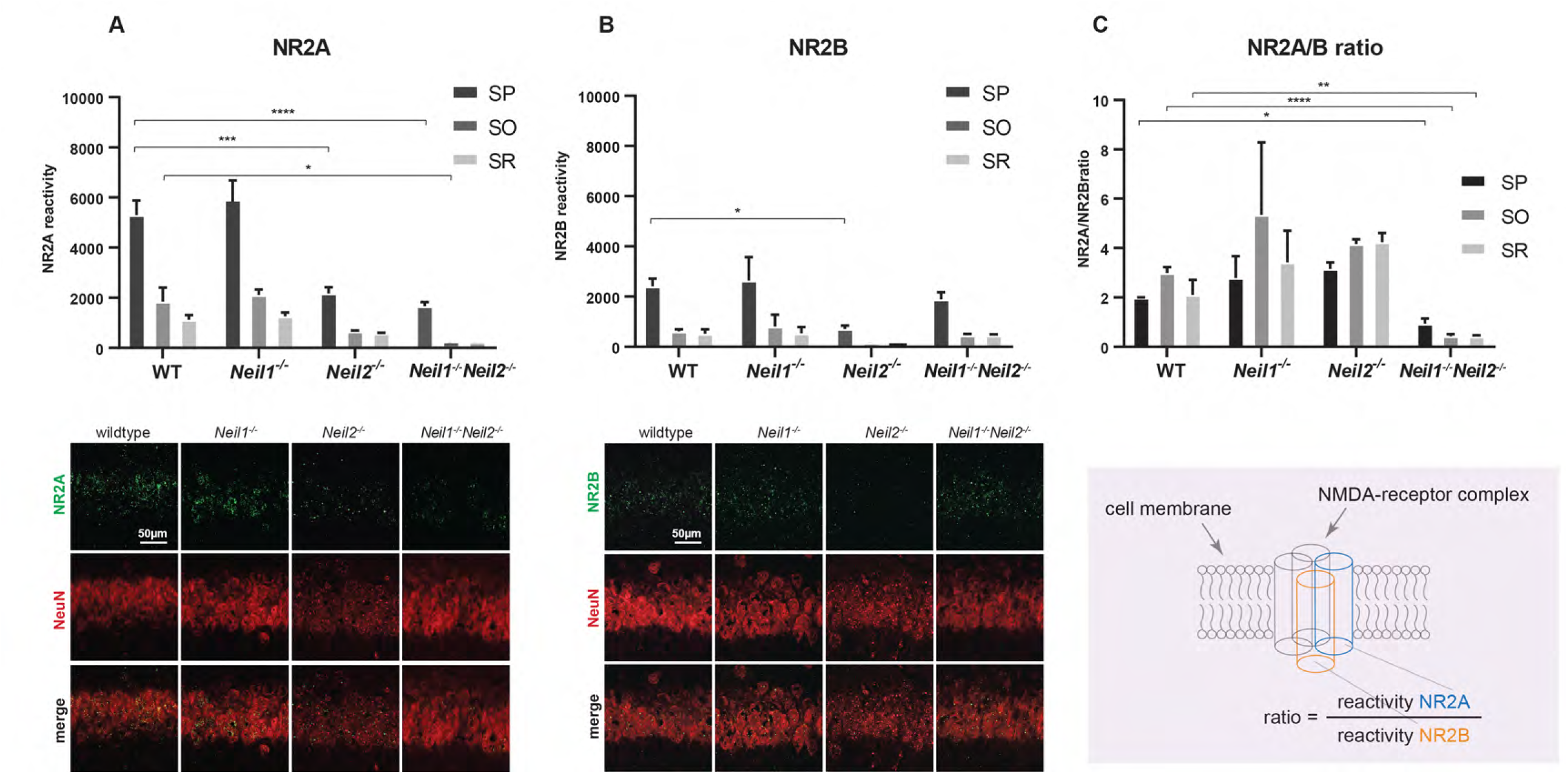
Altered composition of regulatory NMDA-receptor subunits in *Neil1*^*-/-*^*Neil2*^*-/-*^CA1. Immunoreactivity levels of (A) NR2A and (B) NR2B and (C) NR2A vs. NR2B ratio in stratum pyramidale (SP), stratum oriens (SO) and stratum radiatum (SR). Representative images and an illustration of the NMDA-receptor complex are shown. Data are presented as “reactivity levels” based on Imaris spot detection tool (A) and (B) and as ratio of “reactivity levels” (C) + SEM. n = 6 animals (2 slices each), statistics at animal level (see methods). (A) * p = 0.0323, *** p = 0.0003 and **** p < 0.0001 and (B) * p = 0.0354 by two-way ANOVA/Tukey. (C) * p = 0.01, ** p = 0.001, **** p < 0.0001 by multiple t-tests with Holms-Sidak-correction.

Npas4 has been shown to coordinate inhibitory signaling via the GABA-A-receptor, both in vitro and in vivo ^48,49^. We chose to examine specifically the GABA-A-receptor alpha2 subunit (GABRA2) as it is involved in anxiety regulation via distinct intrahippocampal circuits^50^. As for NR2A, GABRA2-reactivity was lowest in *Neil1*^*-/-*^*Neil2*^*-/-*^ mice across SP (Figures 5A and S5A). For *Neil2*^*-/-*^ the difference was close to tendency level (Figure S5A). However, while the expression of NR2A across SO and SR of *Neil2*^*-/-*^ and *Neil1*^*-/-*^*Neil2*^*-/-*^ mice showed a similar tendency to reduction as in SP (Figure 5A), the differences were less conclusive for GABRA2 in SO and SR (Figure S5A).

Postsynaptic density-95 (PSD-95) is an abundant postsynaptic scaffolding protein associated with the NMDA-receptor complex ^51^. In line with reduced absolute levels of NR2A and NR2B in SP of *Neil2*^*-/-*^ and *Neil1*^*-/-*^*Neil2*^*-/-*^ mice (Figure 5A and B), we found a tendency to reduced PSD-95 immunoreactivity in SP in both mutants (Figure S5B).

In sum, these results point to an instability of NMDA-receptor architecture within the postsynaptic compartment in the context of NEIL1/NEIL2 deficiency.

## Discussion

The current study revealed an altered behavioral phenotype in mice deficient in both NEIL1 and NEIL2 DNA glycosylases, shown as increased locomotor activity and reduced anxiety in the open field test and elevated zero maze, and improved learning ability in the Morris water maze test. We have previously reported similar observations in *Ogg1*^*-/-*^*Mutyh*^*-/-*^ mice. However, in these mice, learning was impaired ^16^. Further, we recently demonstrated that mice carrying one deficient allele of *Ogg1* exhibited poorer early-phase learning performance than WT mice using the Barnes maze, and that it was restored when the mice were subjected to oxidative stress by X-ray irradiation ^52^. Inactivation of NEIL3 DNA glycosylase induced an anxiolytic effect and a tendency to impaired learning in mice, however, without increased locomotor activity ^14^. In contrast, overexpression of the repair gene hMTH1, preventing 8-oxoG accumulation in the brain, reduced anxiety in mice without an increase in activity level ^53^. Thus, it appears that DNA glycosylases affect processes involved in behavior and cognition in distinct ways. Canugovi and coworkers previously reported similar learning ability, but defects in short-term spatial memory retention in NEIL1-deficient mice ^23^. Correspondingly, no learning defects were observed in our NEIL1-deficient mice; however, memory was not affected either. A possible explanation to this discrepancy could be that the mice used in the present study were younger (6 months) than the mice tested by Canugovi and colleagues (9-33 months). Further, we have previously shown that *Neil1* mRNA expression increases with age in mouse brains ^29^, suggesting that NEIL1 could be important for cognitive functions at a later stage than we have explored here.

NEIL DNA glycosylases are assumed to be important for genome maintenance by preventing accumulation of oxidative DNA damage. It is therefore reasonable to expect increased levels of oxidative base lesions and possibly mutations when these enzymes are lacking. In line with this, elevated levels of FapyA lesions, but not FapyG or 8-oxoG, were detected in brains from adult (9-22 months) NEIL1 KO mice ^54^. NEIL2 KO mice have also been shown to accumulate oxidized DNA bases in various organs, including brain, but mainly in transcribed regions ^32^. In the present study, accumulation of hippocampal DNA damage was not detected in any of the DNA glycosylase-deficient strains studied and RNA sequencing analysis did not reveal any compensatory upregulation of other DNA glycosylases in hippocampus. Further, only a modest increase in DNA variants in NEIL-deficient hippocampi was found. Although a slightly higher number of variants were detected in the double mutant compared to the single mutants, the number is too small (< 2500 per genome) for the double mutant to be characterized as a hypermutator. Similar observations were made in spleen, liver and kidney of NEIL1/NEIL2-deficient mice, which showed neither increased mutation frequencies nor cancer predisposition under normal physiology ^33^. Further, no global increase in 8-oxoG levels was detected in hippocampus or hypothalamus of mice deficient in both OGG1 and MUTYH DNA glycosylases ^16^. Thus, impaired or reduced global (canonical) repair of oxidized DNA bases in brain regions involved in cognition is not likely to explain the altered behavioral phenotypes observed in DNA glycosylase-deficient mice. Intriguingly, *Neil2*^*-/-*^ and *Neil1*^*-/-*^*Neil2*^*-/-*^ mice showed reduced levels of DNA damage in hippocampus, and we may speculate that this is caused by a putative role of NEIL2 in processes making the chromatin more accessible to strand breaks.

We recently reported that transcriptional changes in the hippocampus of mice lacking OGG1 and MUTYH DNA glycosylases could be an underlying cause of the altered behavioral phenotype observed ^16^. Further, in *Ogg1*^*+/-*^ hippocampus, the expression of three of 35 genes investigated was correlated to spatial learning in the Barnes maze ^52^. Thus, to begin to elucidate the mechanisms behind the behavioral alterations observed in the present study, we looked for changes in the hippocampal transcriptome. We decided to focus on the CA1 subfield of the hippocampal formation, due to its prominent role in spatial information coding and also anxiety-related processes (Bannerman et al., 2014; Buzsáki and Moser, 2013; Jimenez et al., 2018; O’Keefe and Nadel, 1978). RNA sequencing followed by transcriptome analysis revealed that DEGs within the CA1 pyramidal layer of NEIL1/NEIL2-deficient mice referred to genes highly relevant for behavior, synaptic composition and function.

Loss of NEIL2 appears to specifically affect Nr4a orphan receptors, with all three isoforms downregulated in *Neil1*^*-/-*^*Neil2*^*-/-*^ mice, and largely overlapping with *Neil2*^*-/-*^ mice. Consequently, DEGs in *Neil1*^*-/-*^*Neil2*^*-/-*^ were significantly enriched in the nuclear receptor signaling pathway. In CA1, the nuclear receptor signaling pathway is particularly important for regulating excitatory synapse composition ^55^, dopaminergic signaling and, in general, processes of memory formation ^56^. Nr4a1 (Nur77), whose function is enhanced when it forms heterodimers with Nr4a2 (Nurr1) ^57^, interacts reciprocally with (excitatory) NMDA-receptor signaling. It regulates spine density and excitatory synapse distribution especially at distal dendritic compartments ^55^. Further, reduced expression of Nr4a2 has previously been linked to a hyperactive behavior phenotype in mice ^58,59^. This indicates a mechanistically relevant impact of NEIL1 and NEIL2 on these receptors to modulate adaptive behavior.

As another example of NEIL1 and NEIL2 interacting with gene expression relevant for synaptic composition and function, we observed Npas4 to be downregulated both in *Neil1*^*-/-*^ *Neil2*^*-/-*^ and *Neil1*^*-/-*^ animals. Npas4 is prominently involved in regulating the excitatory-inhibitory balance within neural circuits ^44^, with a particular relevance for GABAergic (inhibitory) signaling ^48^. In sum, this suggests that NEIL1 and NEIL2 glycosylases jointly regulate genes relevant for synaptic composition and function, with NEIL2 being prominently involved in nuclear receptor signaling and NEIL1 mainly involved in Npas4-regulation.

With Npas4 being a regulator in excitatory-inhibitory balance and Nr4a receptors interacting directly with the NMDA-receptor, we further examined the expression of the NMDA-receptor in the context of NEIL-deficiency. Here, we focused on the regulatory subunits NR2A and NR2B due to their eminent role in determining the receptor’s electrophysiological properties as well as its relevance in behavior and pathophysiology ^45^. Although total expression levels were decreased in NEIL2- and NEIL1/NEIL2-deficient mice compared to WT, the behavioral phenotype of improved spatial learning is more likely to be explained by the highly reduced NR2A/NR2B ratio, as recent findings suggest a low NR2A/B ratio to be associated with improved memory acquisition performance ^46^ and enhanced LTP ^47^.

While we did not observe differences in LTP, *Neil1*^*-/-*^*Neil2*^*-/-*^ mice displayed electrophysiological changes in the stratum oriens of the hippocampal CA1 subregion as there was a need of higher stimulation strengths to elicit compound action potentials in the Schaffer collaterals of given amplitudes in these mice compared to control mice. This potentially points to a reduced number of afferent fibers in the stratum oriens. Interestingly, recent evidence suggests that the inhibition of heterogeneously tuned excitatory afferent input to CA1 is beneficial for spatial coding ^60^. One could therefore speculate that the reduced excitatory input to CA1 observed in the context of NEIL1/NEIL2 deficiency is sufficient for improved spatial coding at least in the very general spatial learning context of a Morris Water Maze. However, spatial information coding is distinctly a network task involving all hippocampal subfields as well as the entorhinal cortex ^61^. Our study only examines the, albeit very important, CA1 subfield in detail and the behavioral read-out used for this study do not permit conclusions about more refined elements of spatial coding such as pattern completion.

Recently, NEIL1 was identified as a potential reader of oxidized cytosine derivatives, and both NEIL1 and NEIL2 were suggested to potentially cause gene reactivation by an alternative BER pathway for DNA methylation ^62,63^. Furthermore, both proteins were shown to promote substrate turnover by TDG (Thymine-DNA glycosylase) during DNA demethylation^64^. This suggests a role in gene regulation, possibly involving epigenetics that goes beyond canonical repair ^64,65^. In light of this, the behavioral phenotype observed in NEIL1/NEIL2-deficient mice does not seem to be caused by impaired canonical repair of oxidative base lesions. Instead, within CA1, a brain region of utmost importance to numerous cognitive processes, the DNA glycosylases appear to regulate transcription of genes relevant for synaptic function and behavior.

## Conclusion

Here, we identify a cooperative effect of NEIL1 and NEIL2 DNA glycosylases on brain transcription regulation, resulting in a distinct behavioral phenotype with respect to memory formation and anxiety regulation. Our results point to a NEIL1/2 dependent regulation of synaptic factors both at RNA and protein level that is not explained by the enzymes’ function in DNA repair but rather their non-canonical contribution to gene regulation.

## Materials & Methods

### Experimental model and subject details

All experiments were approved by the Norwegian Animal Research Authority and conducted in accordance with the laws and regulations controlling experimental procedures in live animals in Norway and the European Union’s Directive 2010/63/EU.

NEIL1 KO (*Neil1*^*-/-*^), NEIL2 KO (*Neil2*^*-/-*^) and NEIL1/NEIL2 DKO (*Neil1*^*-/-*^*Neil2*^*-/-*^) mouse models generated previously in our lab ^33^ and backcrossed for at least 8 generations onto the C57BL/6N background, were used throughout the study. C57BL/6N mice were included as wild type (WT) controls. Six-month-old male mice were used, unless otherwise stated. The mice were housed and bred in a 12 hour light/dark cycle at the Department of Comparative Medicine, Oslo University Hospital, Rikshospitalet, Norway, or the Comparative Medicine Core Facility, NTNU, Trondheim, Norway, with food and water *ad libitum*. The respective n-value for each experiment is indicated in the main text and/or the figure legends.

### Behavioral studies

Behavioral studies were performed as described previously ^16^ (Figure 1A). Briefly, the Open Field (OF) test ^66^ monitoring general activity, was conducted in an arena measuring L40 cm x W40 cm x H35 cm, where the middle of the arena, L20 cm x W20 cm, was defined as the center area zone. Mice were allowed to explore freely for 45 minutes. The Elevated Zero Maze (EZM) task ^67^ measuring activity and anxiety, was conducted on a circular runway 60 cm above the floor with four alternating open and closed areas. The mice were placed on the maze facing a closed area and allowed 5 minutes for exploration of the apparatus. An open area entry was defined as 85% of the mouse being inside an open area. Learning and memory were monitored using the Morris water maze (MWM) ^68^ Testing was carried out in a white circular pool, 120 cm in diameter and filled 2/3 with white, opaque water (SikaLatex liquid, Sika, Norway) kept at 22 ± 1°C. Using visual cues, the mice learned to find a hidden escape platform, 11 cm in diameter and located at a fixed position 0.5-1.0 cm below the water surface, during repeated daily sessions (days 1-4). The mice were released in the water facing the wall of the pool at four fixed positions in a pseudorandom sequence and given a maximum of 60 sec to locate the hidden platform. Each mouse had eight trials each day in the training period, four in the morning and four in the afternoon. After each block (four trials) the mouse was placed in a heated cage to dry before being returned to the home cage. On days 5 and 12, each mouse was subjected to a single retention trial of 60 sec (probe test) to test spatial memory. During retention trials the escape platform was submerged to the bottom of the pool. A spatial bias for the target quadrant constitutes evidence for spatial memory. During all three behavioral tests, positions of the mice were tracked and stored by using ANY-maze video tracking system (Stoelting, IL, USA). The mice were weighed after the last probe test in the MWM.

### DNA damage analysis

#### HPLC-MS/MS analysis

DNA was isolated from hippocampi (Figure 2A) of WT and NEIL-deficient mice using DNeasy Blood and Tissue kit (Qiagen, cat. no. 80004), according to manufacturer’s protocol. Two µg of genomic DNA was enzymatically hydrolyzed to deoxyribonucleosides by incubation in a mixture of benzonase (Santa Cruz Biotechnology, sc-391121B), nuclease P1 from *Penicillium citrinum* (Sigma, N8630), and alkaline phosphatase from *E. coli* (Sigma Aldrich, P5931) in 10 mM ammonium acetate, pH 6.0, 1 mM magnesium chloride buffer at 40°C for 40 min. Three volume equivalents of ice-cold acetonitrile were added to the reactions after digestion was completed to precipitate proteineous contaminants. Following centrifugation at 16000 × *g* at 4 °C for 40 min, the supernatants were collected in new tubes and dried under vacuum at room temperature. The resulting residues were dissolved in water for HPLC-MS/MS. Chromatographic separation was performed using a Shimadzu Prominence LC-20AD HPLC system with an Ascentis Express C18 2.7 µm 150 x 2.1 mm i.d. column equipped with an Ascentis Express Cartridge Guard Column (Supelco Analytical, Bellefonte, PA, USA) with EXP Titanium Hybrid Ferrule (Optimize Technologies Inc.). For analysis of unmodified nucleosides the following conditions were applied: isocratic flow consisting of 75% A (0.1 % formic acid in water) and 25% B (0.1 % formic acid in methanol) at 0.16 ml/min, 40°C. For analysis of 5-ohC: 0.14 ml/min flow starting with 5% B for 0.5 min, followed with a gradient of 5-45% B for 7.5 min, finishing with re-equilibration with 5% B for 5.5 min. Online mass spectrometry detection was performed using an Applied Biosystems/MDS Sciex API5000 Triple quadrupole mass spectrometer (ABsciex, Toronto, Canada), operating in positive electrospray ionization mode. The deoxyribonucleosides were monitored by multiple reaction monitoring using following mass transitions (m/z): 252.1→136.1 (dA), 228.1 → 112.1 (dC), 268.1→152.1 (dG), 243.1→127.1 (dT) and 244.1→ 128.1 (5-ohdC).

#### Single cell gel electrophoresis (SCGE) / alkaline comet assay

A modified SCGE / alkaline comet assay was performed as previously described ^69^ in a high-throughput format ^70^. Mice were sacrificed and the left hippocampus rapidly dissected using a stereo microscope (Figure 2A). The tissue was immediately placed in ice-cold isotonic solution (Merchant’s buffer; 0.14 M NaCl, 1.47 mM KH_2_PO_4_, 2.7 mM KCl, 8.1 mM Na_2_HPO_4_, 10 mM NaEDTA, pH 7.4, containing EDTA to inhibit cleavage of DNA), mechanically minced to obtain a single cell / nuclei suspension and filtered (100 µm nylon mesh) ^71,72^. The single cell suspensions were counted (Invitrogen Countess™) and diluted to densities appropriate for SCGE (1×10^6^ cells/ml). Cell suspensions were mixed 1:10 with 0.75% Low Melting Point agarose (Gibco BRL 5517US) in PBS, pH 7.4, w/o calcium and magnesium, with 10 mM Na_2_EDTA, to a final agarose concentration of 0.67%. Aliquots of the cell/agarose mixture were instantly added to cold polyester films (GelBond^®^). Solidified gels were immediately immersed in lysis solution (2.5 M NaCl, 0.1 mM EDTA, 10 mM Tris, 1% Sodium Lauryl Sarcosinate, with 1 ml Triton X-100 and 10 ml DMSO per 100 ml solution). After lysis at 4°C overnight, films were washed 1 × 10 min and 1 × 50 min in cold enzyme buffer (40 mM HEPES, 0,1 M KCl, 0,5 mM EDTA, pH 8.0) at 4°C prior to enzyme treatment. To detect oxidative DNA damage, we used the well characterized *E. coli* DNA repair enzyme, Formamidopyrimidine DNA glycosylase (Fpg), as previously described ^69,73–79^. Fpg (1µg/ml) and BSA (0.2 mg/ml) were added to prewarmed enzyme buffer, in which films were immersed and incubated for 1 hour at 37°C. Control films were treated similarly, but with enzyme buffer only (no Fpg added). The Fpg-concentration was optimized based on titration experiments with a photoactivated drug (Ro12-9786) plus cold visible light. After enzyme incubation, films were immersed in cold electrophoresis solution (0.3 M NaOH, 0.1 M EDTA, > pH 13.2) for 5 min + 35 min for unwinding, and electrophoresis was carried out for 25 min at 8-10°C. The voltage potential was 0.80-0.90 V/cm across the stage. Subsequently, films were neutralized in Tris-buffer (0.4 M Tris, pH to 7.5) for 2 × 5 min, rinsed in dH_2_O, fixed in 96% EtOH for 1.5 hours, and dried overnight. Films were rehydrated for 20 min at room temperature in TE-buffer pH 7.5, containing 10,000 × diluted SYBRGold stain (Molecular Probes), under gentle shaking. The films were rinsed in dH_2_O and covered with large coverslips (80 × 120 mm, thickness no.1, VWR International AS, Oslo, Norway). Imaging was performed with an epi-fluorescence microscope (Olympus BX51). Semi-automated scoring of 50 comet tails per gel was done with “Comet assay IV” software (Perceptive Instruments Ltd, UK). The median comet tail intensity per sample (50 comets × 3 replicate gels scored) was used to calculate the mean values per genotype. Net Fpg-sensitive sites were calculated by subtracting the median comet tail intensity for samples incubated without Fpg from those treated with Fpg.

#### PCR-based DNA damage detection

Hippocampal DNA damage levels were quantified by using a restriction enzyme-based qPCR method as described previously ^80^ Briefly, DNA damage in a TaqI-sensitive restriction site will result in altered cutting frequency of the DNA, which ultimately will affect PCR amplification of a target sequence spanning the restriction site. Total genomic DNA was isolated from hippocampus (Figure 2A) using the DNeasy Blood and Tissue kit according to manufacturer’s protocol (Qiagen, cat. no. 80004). 30 ng of DNA was subjected to *Taq*^α^I restriction enzyme digestion followed by qPCR amplification of a target sequence in the *Gapdh* gene. Relative amounts of PCR products, reflecting the level of damage in each sample, were calculated by using the comparative ΔCT method. Primers: *Gapdh* forward, 5’cttcaacagcaactcccact and reverse, 5’aaaagtcaggtttcccatcc.

### DNA mutation analysis

#### Whole-genome deep sequencing

For each genotype, hippocampal genomic DNA from four naïve mice was isolated using DNeasy Blood and Tissue Kit (Qiagen, cat. no. 80004), pooled and sent to BGI Tech Solutions, Hong Kong, for whole genome sequencing, including library construction and HighSeq4000 sequencing.

#### Identification of strain-dependent genetic variations

We identified SNPs and insertions/deletions (InDels) individually for mutant and WT samples. Specifically, adapter sequence in the raw data was removed, and low-quality reads which had too many Ns (>10%) or low-quality score (<5) was discarded. The remaining reads were aligned to the mouse reference sequence (mm10) using the Burrows-Wheeler Aligner (BWA) ^81^. The alignment information was stored in BAM format files, which was further processed by fixing mate-pair information, adding read group information and marking duplicate reads caused by polymerase chain reaction artefacts. The variant calling steps included SNPs detected by SOAPsnp ^82^ and small InDels detected by Samtools/GATK ^83^. In GATK, the caller *UnifiedGenotyper* was used with the parameters *stand_call_conf* set to 50 and *stand_emit_conf* set to 10. Hard filtering was applied to get variant results of higher confidence. To identify strain-dependent genetic variation – i.e. variants inherited from the 129 strain and not completely lost through back-crossing with the C57BL/6N strain – SNP and InDel data were loaded into the genome browser *SeqMonk* (http://www.bioinformatics.babraham.ac.uk/projects/seqmonk/) for further inspection. We defined 129-specific regions as having more than 50 detected SNPs or InDels per 600 kB bases and used this as criterion in the “Read Count Quantitation using all Reads” probe extraction method in *SeqMonk*. Individual regions satisfying this criterion were extracted and consecutive regions within the genome were joined to form the final 129-dependent regions. We confirmed enrichment of 129-dependent genetic variants within each region by identifying the SNPs that were present in dbSNP (build 137) and counting the number of times the SNP genotype matched the annotated 129 (129P2/OlaHsd, 129S1/SvImJ or 129S5SvEv strains) or black 6 (C57BL/6NJ strain) genotypes.

#### Identification of mutations in NEIL-deficient hippocampi

Reads were filtered and aligned to the mouse genome as described above, and alignments were preprocessed according to GATK Best Practices recommendations ^84^ using GATK version 3.5, including local realignment around InDels and recalibration of quality scores. For calling we used the MUTECT2 variant caller ^85^, with KOs as case and WT as control. Briefly, MUTECT2 identifies variants that are present in the case sample but are absent in the control sample and where the difference is unlikely due to sequencing errors. We used MUTECT2 default parameters, which include rejecting candidates that in the control sample have (i) supporting reads numbering ≥ 2 or constituting ≥ 3% of the total reads (i.e. < 34 total reads) and (ii) their quality scores sums to > 20. We used snpEff ^86^ and SnpSift ^87^ to annotate all SNPs and InDels found and discarded SNPs and InDels overlapping the 129-specific intervals for each sample.

### Electrophysiology

#### Slice/Sample preparation

Adult (4-month-old) WT and *Neil1*^*-/-*^*Neil2*^*-/-*^ male and female mice were sacrificed with Suprane (Baxter) and the brains removed. Transverse slices (400 μm) were cut from the middle and dorsal portion of each hippocampus with a vibroslicer in artificial cerebrospinal fluid (ACSF, 4°C, bubbled with 95% O_2_ -5% CO_2_) containing (in mM): 124 NaCl, 2 KCl, 1.25 KH_2_PO_4_, 2 MgSO_4_, 1 CaCl_2_, 26 NaHCO_3_ and 12 glucose. Slices were placed in an interface chamber exposed to humidified gas at 28-32°C and perfused with ACSF (pH 7.3) containing 2 mM CaCl_2_ for at least one hour prior to the experiments. In some of the experiments, DL-2-amino-5-phosphopentanoic acid (AP5, 50uM; Sigma-Aldrich, Oslo, Norway) was added to the ACSF in order to block NMDA receptor-mediated synaptic plasticity.

#### Synaptic transmission, synaptic excitability and paired-pulse facilitation

Orthodromic synaptic stimuli (<300 µA, 0.1 Hz) were delivered through tungsten electrodes placed in either stratum radiatum or stratum oriens of the hippocampal CA1 region. The presynaptic volley and the field excitatory postsynaptic potential (fEPSP) were recorded by a glass electrode (filled with ACSF) placed in the corresponding synaptic layer while another electrode placed in the pyramidal cell body layer (stratum pyramidale) monitored the population spike. Following a period of at least 20 min with stable responses, we stimulated the afferent fibers with increasing strength (increasing the stimulus duration in steps of 10 µs from 0 to 90 µs, five consecutive stimulations at each step). Prior to the input/output (I/O), the strength was adjusted so that a population spike appeared in response to 40, 50, or 60 µs in order to define the stimulation/response range. A similar approach was used to elicit paired-pulse responses (50 ms interstimulus interval, the two stimuli being equal in strength). To assess synaptic transmission, we measured the amplitudes of the presynaptic volley and the fEPSP at the different stimulation strengths. During the analysis, care was taken to use extrapolated measurements which were within the apparently linear part of the I/O curves. Values from individual experiments outside the linear part of the I/O curves (prevolley vs. fEPSP) were omitted when pooling the data. The population spike amplitude was measured as distance between the maximal population spike amplitude and a line joining the maximum pre- and postspike fEPSP positivities. In order to pool data from the paired-pulse experiments we selected responses to stimulation strength just below the threshold for eliciting a population spike on the second fEPSP.

#### Long-term potentiation (LTP) of synaptic transmission

Orthodromic synaptic stimuli (50 μs, < 300 μA) were delivered alternately through two tungsten electrodes, one situated in the stratum radiatum and another in the stratum oriens of the hippocampal CA1 region. Extracellular synaptic responses were monitored by two glass electrodes (filled with ACSF) placed in the corresponding synaptic layers. After obtaining a stable synaptic response in both pathways (0.1 Hz stimulation) for at least 10-15 minutes, one of the pathways was tetanized (a single 100 Hz tetanization for 1 sec, repeated four times at 5 min intervals). As standardization, the stimulation strength used for tetanization was just above threshold for generation of a population spike in response to a single test shock. Synaptic efficacy was assessed by measuring the slope of the fEPSP in the middle third of its rising phase. Six consecutive responses (1 min) were averaged and normalized to the mean value recorded 1-4 min prior to tetanization. Data were pooled across animals of the same experimental group and pathway and are presented as mean ± SEM. *Frequency facilitation and delayed response enhancement*. In order to compare other forms of short-term synaptic plasticity in the two genotypes, we activated either the radiatum or the oriens pathway at 20 Hz for one minute following the presence of stable synaptic responses (0.1 Hz stimulation) for at least 10-15 minutes. The synaptic strength was assessed by measuring the maximal slope of the rising phase (V/s) of the fEPSPs, and normalizing the value of each response to the mean value recorded one minute prior to the switch to a higher stimulation frequency. To compare the processes developing during the 20 Hz stimulation train, we compared the magnitude of the frequency facilitation and delayed response enhancement (DRE) in the two genotypes. Three time points were also used to characterize the time courses of the different phases during the 20 Hz stimulation, i.e. i) the time needed to reach the maximal magnitude of the initial frequency facilitation; ii) the transition point where the minimal value of the response during the subsequent decay period was observed; and iii) the time point of the maximal value reached during the DRE phase.

### Laser capture microdissection and RNA sequence analysis

#### Tissue processing

Mouse brains (n=3 mice/genotype) were isolated without prior perfusion within 150+/-30s, mounted onto cryostat sockets (Leica CM3050S, Nussloch, Germany) and frozen in liquid nitrogen immediately. Subsequently, samples were stored in liquid nitrogen until further processing (max. 1hr). Each brain was completely processed according to the workflow described below the same day. Brains were cut in coronal orientation at a thickness of 8µm using a cryostat (Leica CM3050S, Nussloch, Germany), starting from the onset of the hippocampal formation until the end. The Allen Mouse Brain Atlas (Allen Brain Institute) was used as a reference. 20-30 laser dissection membrane slides (Molecular Machines and Industries GmbH, Eching, Germany) with 5-7 brain slices each were collected from each brain. All slides were subsequently used for tissue collection to avoid a collection bias alongside the rostro-caudal axis.

#### Tissue collection

CA1 dissectates were collected using a laser dissection microscope (Molecular Machines and Industries, CellCut on Olympus IX71, Eching, Germany). Only dorsal hippocampus was included in the tissue collection described in the following. The hippocampal CA1 area was identified using the Allen Mouse Brain Atlas as a reference. For each slice, the whole CA1 area was defined manually (see Figure 4A). We collected 20 CA1 dissectates in one isolation cap (Molecular Machines and Industries GmbH, Eching, Germany) before adding RLT lysis buffer (AllPrep Kit, Qiagen, Hilden, Germany). Samples from one individual were collected the same day. Samples were vortexed shortly and frozen at -80°C until further processing.

#### RNA extraction

RNA was isolated using the RNeasy Mini Kit (Qiagen, Hilden, Germany) according to the manufacturer’s instructions. Samples were briefly vortexed before starting the RNA isolation steps; no additional tissue lysis procedure was performed. RNA samples yielded >280ng of RNA (>5.6ng/µl in a total eluate of 50µl) with a RIN value of generally > 7 as determined by Bioanalyzer (Agilent Technologies, Santa Clara, USA).

#### RNAseq

Whole-transcriptome sequencing was done by BGI Group (BGI Genomics Co., Ltd., Hong Kong, China). In brief, the steps were as follows: a) quality control using the Agilent 2100 Bio analyzer (Agilent RNA 6000 Nano Kit, Agilent Technologies, Santa Clara, CA, USA), b) purification of poly-A containing mRNA by poly-T oligo-attached magnetic beads, c) mRNA-fragmentation (divalent cations, high temperature), d) reverse transcription into cDNA, e) Qubit quantification (Thermo Fisher Scientific, Waltham, MA, USA), f) library construction by making a single strand DNA circle (see library construction quality control results Figure S4 H), g) rolling circle replication creating DNA nanoballs (more fluorescent signal during sequencing), h) reading of pair-end reads of 100bp via BGISEQ-500 platform ^88^.

#### RNAseq bioinformatic preprocessing

Bioinformatic processing was in part done by BGI (BGI Genomics Co., Ltd., Hong Kong, China) using the following workflow: a) filtering of low quality reads using the software SOAPnuke ^89^, b) genome mapping with HISAT software ^90^, c) transcript reconstruction using StringTie ^91^ and reference comparison with Cuffcompare ^92^, d) prediction of coding potential with CPC ^93^, e) SNP and INDEL detection with GATK ^83^, e) reference mapping with Bowtie2 ^94^, f) calculation of gene expression levels using the RSEM software package ^95^, g) hierarchical clustering with hclust in R.

#### Analysis of differential expression

We created the count matrix of integer values and the metadata matrix based on the sequencing results from BGI (un-normalized estimated counts). We next performed a DESeq2 differential gene expression analysis (R version “Dark and Stormy Night” for Windows ^96^ comparing between the different genotype groups (n = 3 mice per group; *Neil1*^*-/-*^, *Neil2*^*-/-*^, *Neil1*^*-/-*^*Neil2*^*-/*-^, WT). Low counts were filtered using rowSums function. The alpha parameter of the results function was set to 0.05 and adjusted p-value was calculated using Benjamini-Hochberg correction for multiple testing (see all DEGs Figure S4, D-F). We performed this analysis after prior exploratory data analysis based on the BGI-results, which showed two samples to be clear outliers from the remaining replicates (PCA-plot, Figure S4 G). These two outliers (red arrows) were excluded from the group comparison analysis. For the over-representation analysis the online version of PANTHER Classification System release 15.0 was used ^97^. We chose a Binomial/FDR multiple testing correction, both for the gene ontology biological processes term enrichment analysis and the reactome-pathway enrichment analysis (Reactom version 73 2020/06/11) ^98^. All DEGs log2fold>0.3 (fold>1.23) were included in the enrichment analysis (Figure S4 D-F).

### Immunohistochemistry

#### Mouse perfusion

Mice were anesthetized and killed using first isoflurane (Baxter, Oslo, Norway) and subsequently an intraperitoneal, weight-adapted overdose of pentobarbital (>200mg/kg bodyweight). Intracardial perfusion was done with 0.9% saline (B.Braun, Melsungen, Germany) and 4% paraformaldehyde in phosphate-buffered saline (PBS). Brains were put into 4% paraformaldehyde/PBS solution for a minimum of 48hrs at 4°C for fixation. We sectioned brains at a thickness of 30µm using a cryostat (Leica CM3050S, Nussloch, Germany) starting at a medio-lateral depth of 900µm and continuing until the end of the tissue block. Slices were then stored at 4°C in a PBS-solution containing 0.05% of Proclin (Merck, Darmstadt, Germany) until further processing.

#### Antibody treatment

Heat-mediated antigen retrieval was performed for 3 min at 99°C in a 40mM trisodium citrate (Merck, Darmstadt, Germany) solution, pH 6.0. Specimens were then left to cool down to room temperature inside this solution for another 27min (30min total exposure). 5% normal goat serum/bovine serum albumin served as a blocking agent against unspecific binding. We then immediately incubated in primary antibody solution (GABRA2 rabbit polyclonal, Cat.No.224103, Synaptic Systems, Goettingen, Germany; NR2A rabbit polyclonal, Cat.No. AGC-002, alomone labs, Jerusalem, Israel; NR2B rabbit polyclonal, Cat.No. AGC-003, alomone labs, Jerusalem, Israel; NeuN mouse monoclonal, A40/MAB377, Cat.No. 636574, ThermoFisher, Waltham, MA, USA; PSD-95 rabbit polyclonal, Cat.No. PA585769, ThermoFisher, Waltham, MA, USA) over night at 4°C (exception: PSD-95 2 hours at room temperature, no antigen retrieval step) under constant shaking at 15 oscillations/min. We generally used Alexa Flour dyes (A488, A555, A647) as secondary antibodies (Thermo Fisher Scientific, Waltham, MA, USA) and DAPI (Merck, Darmstadt, Germany) as a non-specific nuclear counter stain.

#### Confocal microscopy

Imaging of stained slices was done using a Zeiss LSM880 confocal microscope. For synaptic markers, a Plan-Apochromat 40x/1.4 Oil DIC M27 objective (Carl Zeiss, Jena, Germany) was used. An imaging square of 700×700µm (x/y 2000pixels of 0.35µm each) and a z-interval of 0.5µm was applied. A proximal and a distal imaging square was set within the CA1 region based on NeuN and DAPI stainings as an anatomical orientation (center of proximal square set at 0.25x total CA1 length measured from proximal end, center of distal square at 0.75xtotal CA1 length). Results displayed are averaged across proximal and distal squares. For each animal, one medial and one lateral brain slice were analyzed, amounting to a total of 4 imaging squares (2 proximal, 2 distal). With respect to the nested data problem (Aarts et al, NatNeurosci 2014), all statistical analysis was done at an animal level (1 animal = 1 statistical unit). On a sideline, we observed a consistently different NeuN-signal in NEIL2-deficient mice. Tissue quality, immunostaining protocol parameters, background signal and DAPI-counter-staining efficiency was identical in these samples compared to the other genotypes, so that a genotype-specific NeuN-signal appears possible and, while beyond the scope of this manuscript, warrants further investigation.

#### Quantification of immunoreactivity

Imaris 9.3 (Bitplane, Zurich, Switzerland) was used to quantify immunoreactivity. We first created a 3D reconstruction of the whole z-plane dataset. The strata pyramidale/oriens/radiatum were identified as regions of interest using the “surface” tool in Imaris and copied to every z-plane accordingly. Based on the surface selection, a 3D-frame was created and the parameter of interest “masked” according to this frame. The Imaris “Spots Wizard” function was used to identify areas of synaptic reactivity within this masked channel (1µm spot diameter, background subtraction applied according to local contrast). We then conducted a pilot-experiment for every immunohistochemical marker used, involving typically 4 different images. This was to make sure a biologically relevant signal is captured by the software. Based on this pilot experiment, a selection criterion based on the “quality” (see bitplane.com/imaris) filter in Imaris was defined and kept the same throughout the analysis. Imaris automated background subtraction was done for every specimen analyzed to account for intensity variations despite identical immunohistochemistry and confocal parameters. 2/3 of the region of interest had to be intact (i.e. not damaged by tissue cracks, covered by imaging artefacts etc.) in order to be included in the analysis. One exclusion was made based on this criterion (NR2A immunostaining, wildtype animal). No outlier correction was performed before statistical analysis. No image processing was performed prior to Imaris quantification (raw image data, Zeiss “lsm” file format). Figure 5 shows a background-filtered signal indicating the approximate portion detected by the spot-detection tool.

### Quantification and statistical analysis

In figures 1B-I, 2B, S1 and S2 statistical evaluation was done by one-way ANOVA followed by Tukey’s multiple comparisons test using GraphPad Prism version 6.07 for Windows (GraphPad Software, La Jolla California USA, www.graphpad.com). Data in figures 1J and 1K-L were analyzed by non-parametric pair wise Wilcoxon rank-sum test, Holm-adjusted, and post-hoc family-wise multiple comparison of means (Tukey Honest Significant Difference), respectively, using R version 3.0.2. In figure 2C, the data were analyzed by two-way ANOVA, followed by Sidak’s multiple comparisons test using GraphPad Prism version 8.4. In figures 2D-G the MuTect2 (Genome Analysis Toolkit (GATK) toolset) software was used. In figures 3 and S3, statistical evaluation was done by linear mixed model analysis, using SAS 9.2 (SAS Institute Inc.). For statistical analysis of RNA seq data (figures 4 and S4), we refer to the Methods section. Data in figure S5B was analyzed by one-way ANOVA and in 5A-B and S5A by two-way ANOVA using GraphPad Prism version 8.4. Tukey’s multiple comparisons test was performed in all cases. In figure 5C, multiple t-tests with Holm-Sidak correction, was applied. A p-value < 0.05 was considered significant.

### Data availability

Further information and requests for resources and reagents should be directed to and will be fulfilled by the Lead Contact, Magnar Bjørås. The RNA sequence data discussed in this publication have been deposited in NCBI’s Gene Expression Ominbus ^99^ and are accessible through GEO Series accession number GSE160621 (https://www.ncbi.nlm.nih.gov/geo/query/acc.cgi?acc=GSE160621). The DNA sequence data have been deposited in European Nucleotide Archive (ENA), accession number PRJEB31108.

## Acknowledgements

We thank our colleague and friend Øivind Hvalby^✝^ who actively contributed to the electrophysiological data collection and analysis in this study but sadly passed away in 2015. We thank Atle van Beelen Granlund for his help in setting up the laser capture microdissection experiment. The LC-MS/MS analysis of DNA modifications was performed by the Proteomics and Modomics Experimental Core Facility (PROMEC), Norwegian University of Science and Technology (NTNU).

This work was sponsored by the Research Council of Norway [287911, 240529]; the South-Eastern Norway Regional Health Authority [46060921, 2017117]; and partly supported by the Research Council of Norway through its Centers of Excellence funding scheme [223268/F50]. PROMEC is funded by the Faculty of Medicine and Health Sciences at NTNU and the Central Norway Regional Health Authority.

## Author contributions

G.A.H. and L.L generated the NEIL1 KO mouse model in collaboration with the Norwegian Transgenic Center. R.S. generated the NEIL2 KO mouse model and backcrossed and bred the mice. N.K. backcrossed and bred the mice in Trondheim. V.R., O.M. and M.B. designed behavioral experiments. V.M., R.S., S.V. and M.D.B. conducted behavioral experiments. A.D.R. analyzed behavioral results, including statistics. A.K. performed HPLC-MS/MS. A.K.O. and K.B.G. performed alkaline comet assay. K.S. performed PCR based DNA damage assay. P.S. analyzed DNA sequencing data. V.J. designed and conducted electrophysiological studies and analyzed the data obtained. N.K. designed the laser capture microdissection experiment. N.K., S.B.S. and M.S.F. performed the laser capture microdissection experiment. N.K. isolated RNA from laser-dissected samples. M.S.F. performed the immunostaining experiments. N.K. did the confocal imaging and subsequent Imaris-analysis as well as statistical analysis. W.W. performed qPCR. A.M.B. analyzed the RNAseq data acquired from BGI. A.M.B. and N.K. performed the gene ontology and pathway analysis of RNAseq data. N.K. did the single-gene QuickGo thematic analysis. N.K. and G.A.H. layouted, revised and structured all figures based on single plots and figures made by other authors. G.A.H., V.R., N.K, K.S. and M.B. wrote the manuscript with input from the other authors.

