## Supplementary Material for "NEIL1 and NEIL2 DNA glycosylases regulate anxiety and learning in a cooperative manner"

Medicine,

[†] The first three authors should be regarded as Joint First Authors.

[§] The last two authors should be regarded as Joint Last Authors.

**Figure S1**

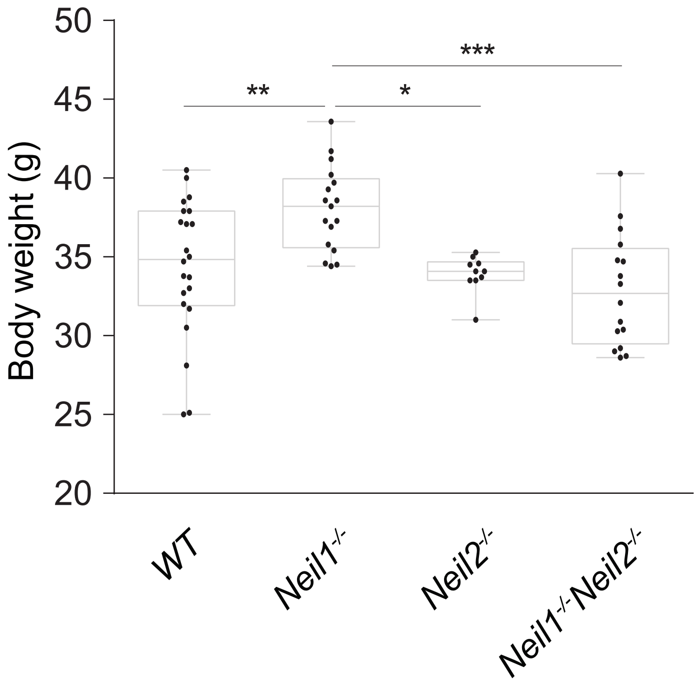


**Figure S1. Increased body weight in NEIL1-deficient mice.**

The mice were weighed after the behavioral studies were completed. Data are shown in full, with overlaid boxplots representing the medians and interquartile ranges (IQR), Whiskers indicate min/max values. n = 22 WT, 17 *Neil1*$^{-/-}$, 10 *Neil2*$^{-/-}$ and 16 *Neil1*$^{-/-}$*Neil2*$^{-/-}$ mice. * $p < 0.0185$, ** $p < 0.0073$, *** $p < 0.0003$ by one-way ANOVA/Tukey.

**Figure S2**

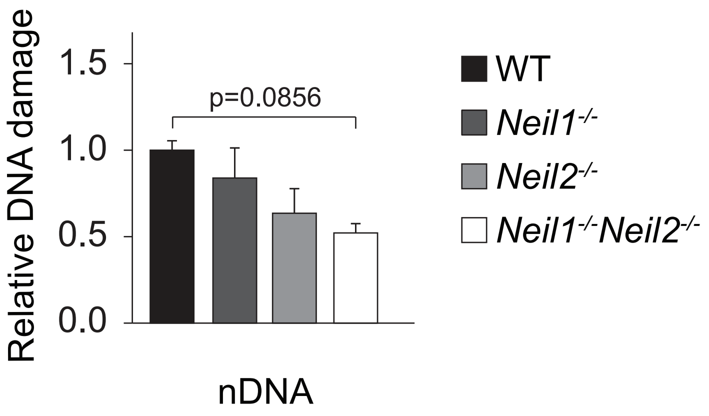


**Figure S2. Tendency to reduced DNA damage level in NEIL1/NEIL2 -deficient mice.** Hippocampal DNA damage was measured by using a restriction enzyme-based qPCR method detecting damage in the *Gapdh* gene (1). Data are shown as mean + SEM. n = 3 WT, 3 *Neil1*$^{-/-}$, 4 *Neil2*$^{-/-}$ and 4 *Neil1*$^{-/-}$*Neil2*$^{-/-}$ mice. p-value by one-way ANOVA/Tukey.

**Figure S3**

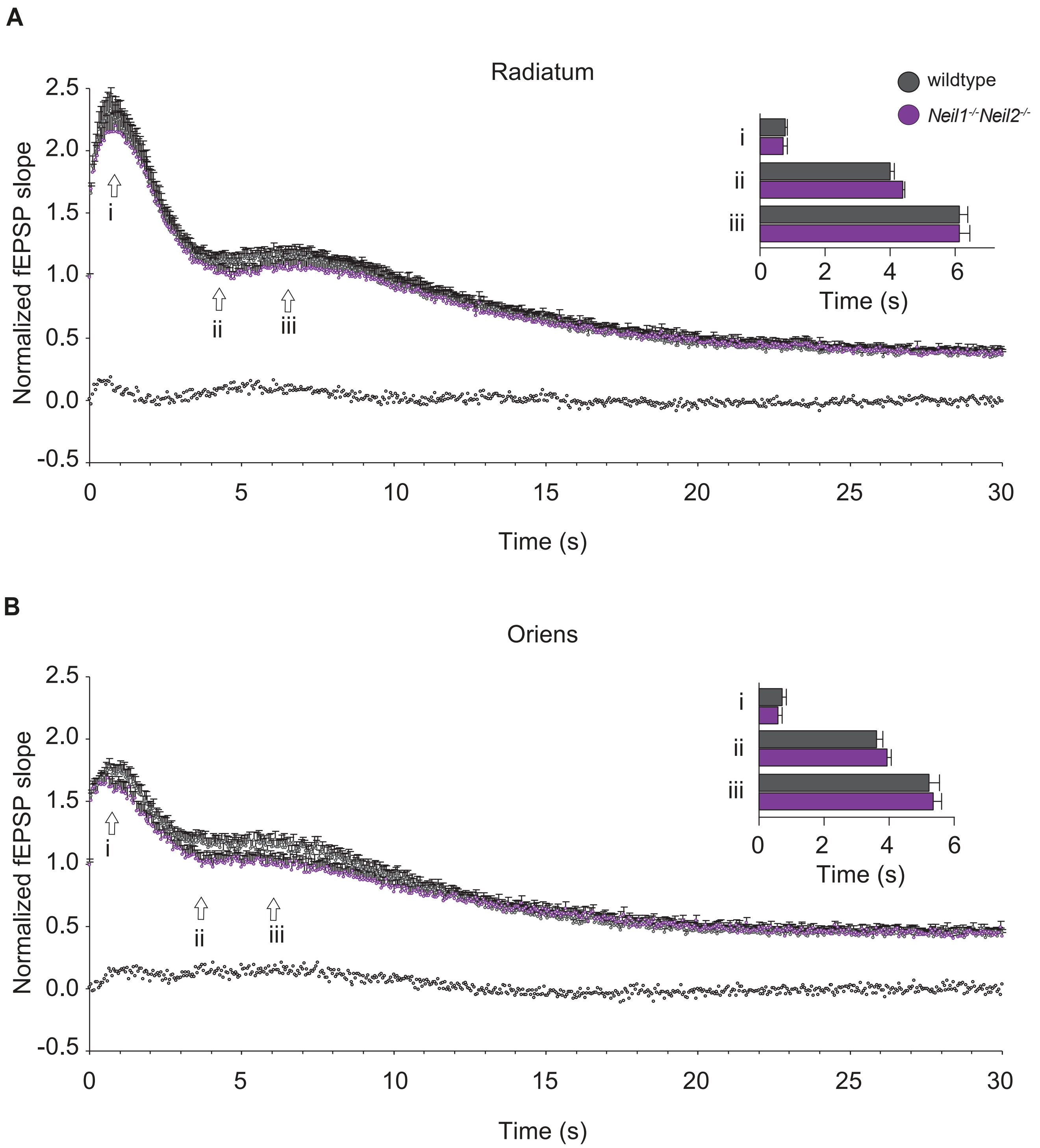


**Figure S3. Normal delayed response enhancement at hippocampal CA3 to CA1 synapses in NEIL1/NEIL2-deficient mice.** Normalized and pooled fEPSP slope measurements during the initial 30 s of 20 Hz stimulation trains in stratum radiatum (SR) and stratum oriens (SO). Arrows indicate approximate positions for: (i) the time point to the maximum magnitude of the initial frequency facilitation, (ii) time to the transition point between the initial frequency facilitation and the delayed response enhancement (DRE), and (iii) time needed to reach the peak of the DRE. Black horizontal bar along the abscissa indicate $p < 0.05$ when comparing the genotypes. The histograms show comparisons between time points i, ii, and iii in WT and *Neil1*$^{-/-}$*Neil2*$^{-/-}$ mice. Data are shown as mean + SEM. SR, n = 17 WT and 18 *Neil1*$^{-/-}$*Neil2*$^{-/-}$; SO, n = 15 WT and 17 *Neil1*$^{-/-}$*Neil2*$^{-/-}$ mice.

### Figure S4

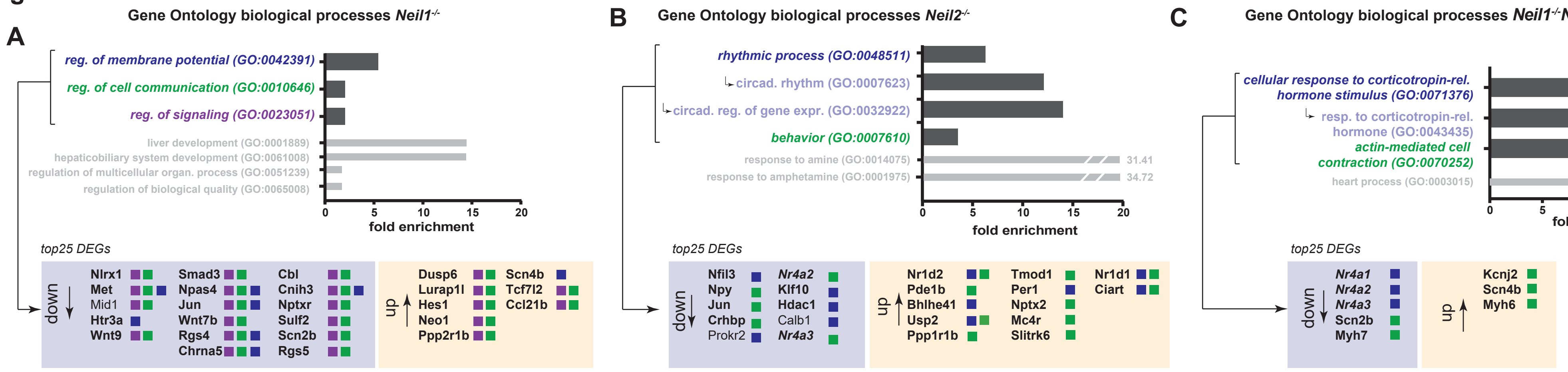


**D** DEGs *Neil1-/-*

| log2fold | Symbol | log2fold | Symbol | log2fold | Symbol |
|---|---|---|---|---|---|
| -6.33 | *Gm45988* | -0.37 | *Pkm* | 3.19 | *Gm29801* |
| -2.30 | *Nlrx1* | -0.36 | *Tspan3* | 3.90 | *Gm16867* |
| -2.21 | *Clmp* | -0.35 | *Sprn* | 4.27 | *Gm29804* |
| -2.17 | *Met* | -0.35 | *Lgi1* | 5.86 | *Mpzl2* |
| -1.96 | *Mid1* | -0.35 | *Fam210b* | 6.05 | *Treh* |
| -1.55 | *Htr3a* | -0.33 | *Npcd* | 6.11 | *Xntrpc* |
| -1.50 | *Wnt9a* | 0.34 | *Dnajb5* | 6.51 | *Gm13302* |
| -1.43 | *Npbwr1* | 0.34 | *Htr1a* | 8.81 | *Gm2427* |
| -1.41 | *Car12* | 0.36 | *Glt8d2* | 9.99 | *LOC100041599* |
| -1.39 | *H2ac24* | 0.37 | *Gpt2* | 10.71 | *Ccl21b* |
| -1.36 | *Cryab* | 0.39 | *Tra2a* | | |
| -1.36 | *Iho1* | 0.40 | *Rplp1* | | |
| -1.07 | *Pcdha6* | 0.40 | *Sesn1* | | |
| -1.03 | *Smad3* | 0.40 | *Pid1* | | |
| -1.03 | *Npas4* | 0.40 | *Extl2* | | |
| -0.99 | *Jun* | 0.43 | *Ier5* | | |
| -0.91 | *Thsd4* | 0.44 | *Rab8b* | | |
| -0.86 | *Cd276* | 0.44 | *Bhlhe40* | | |
| -0.84 | *Prokr2* | 0.46 | *Dusp6* | | |
| -0.83 | *Wnt7b* | 0.47 | *Kdm7a* | | |
| -0.80 | *Rgs4* | 0.50 | *Ormdl3* | | |
| -0.72 | *Cbl* | 0.51 | *Lurap1l* | | |
| -0.68 | *Cnih3* | 0.54 | *Srsf10* | | |
| -0.67 | *Sst* | 0.55 | *Phldb1* | | |
| -0.64 | *Zfp46* | 0.65 | *Pcdha12* | | |
| -0.62 | *Npy* | 0.66 | *Hes1* | | |
| -0.62 | *Usp2* | 0.68 | *Neo1* | | |
| -0.60 | *Nptxr* | 0.72 | *Pcdhga6* | | |
| -0.57 | *Sulf2* | 0.72 | *Lhfp* | | |
| -0.54 | *Rhou* | 0.72 | *Dcps* | | |
| -0.54 | *Gm40363* | 0.78 | *Ppp2r1b* | | |
| -0.53 | *Scn2b* | 0.79 | *Slc37a4* | | |
| -0.52 | *Rgs5* | 0.85 | *Entpd4b* | | |
| -0.52 | *Ndn* | 0.86 | *Ift46* | | |
| -0.50 | *Chrna5* | 0.91 | *Snx22* | | |
| -0.50 | *Kcna4* | 0.94 | *Scn4b* | | |
| -0.48 | *St6gal2* | 1.06 | *Tmem254b* | | |
| -0.45 | *Tcf12* | 1.14 | *Pcdhb4* | | |
| -0.43 | *Wdr6* | 1.16 | *Txnip* | | |
| -0.43 | *Bace1* | 1.18 | *Tmem254a* | | |
| -0.40 | *Dnaja4* | 1.39 | *Lrrc1* | | |
| -0.40 | *Coro2b* | 1.76 | *H2ac23* | | |
| -0.40 | *Sema7a* | 1.90 | *Fxyd2* | | |
| -0.39 | *AI593442* | 2.13 | *Tcf7l2* | | |

**All DEGs: log2fold >0.3, $p_{adj}$<0.05**

**E** DEGs *Neil2-/-*

| | | | | | | | |
|---|---|---|---|---|---|---|---|
| -3.05 | *Sftpc* | -0.50 | *Bmp1* | 0.34 | *Hgsnat* | 0.68 | *Acss1* |
| -2.63 | *Chst9* | -0.50 | *Hspa5* | 0.35 | *C1qtnf4* | 0.72 | *Cirbp* |
| -2.18 | *Gm43305* | -0.48 | *Gm40363* | 0.36 | *Fdft1* | 0.78 | *Mc4r* |
| -2.10 | *Mid1* | -0.47 | *Pdia6* | 0.37 | *Ccdc85b* | 0.80 | *St8sia6* |
| -2.05 | *Xlr3b* | -0.46 | *Rbm28* | 0.37 | *Pde1b* | 0.87 | *Il16* |
| -1.86 | *Dcn* | -0.46 | *Hdac1* | 0.38 | *Slc45a1* | 0.95 | *Dse* |
| -1.83 | *Npbwr1* | -0.46 | *Alcam* | 0.39 | *Rasl11b* | 0.97 | *Rgcc* |
| -1.49 | *Cort* | -0.45 | *Rasl10a* | 0.39 | *Pcdhga2* | 1.02 | *Tmem254b* |
| -1.41 | *Rprm* | -0.45 | *Tra2a* | 0.39 | *Luzp2* | 1.04 | *Slitrk6* |
| -1.31 | *Gfra2* | -0.43 | *Rngtt* | 0.39 | *Vstm2l* | 1.09 | *Nr1d1* |
| -1.28 | *Myh7* | -0.43 | *H2bc4* | 0.40 | *Ip6k2* | 1.16 | *Nppc* |
| -1.23 | *Slc17a6* | -0.42 | *Psg16* | 0.40 | *Rab36* | 1.21 | *Tmem254a* |
| -1.10 | *Esd* | -0.42 | *Calb1* | 0.41 | *Ppif* | 1.24 | *Klhl33* |
| -1.03 | *Kcnc4* | -0.39 | *Nptx1* | 0.41 | *Bhlhe41* | 1.40 | *Gm14685* |
| -1.03 | *Pcdha1* | -0.36 | *Rexo4* | 0.42 | *2900026A02Rik* | 1.71 | *Ciart* |
| -1.00 | *Slc39a2* | -0.36 | *Nr4a3* | 0.42 | *Usp2* | 1.96 | *LOC108168952* |
| -1.00 | *Gm14434* | -0.34 | *Hsp90b1* | 0.42 | *Mcur1* | 1.96 | *LOC108168680* |
| -0.97 | *Nfil3* | -0.33 | *Pdia3* | 0.42 | *Mex3b* | 2.84 | *Gm29801* |
| -0.96 | *Crlf1* | -0.33 | *Dmtf1* | 0.43 | *Tlnrd1* | 4.26 | *Gm29804* |
| -0.92 | *Cldn5* | -0.33 | *Lman2l* | 0.43 | *Kdm4b* | 4.47 | *Myh6* |
| -0.92 | *Lrrtm3* | -0.33 | *Ptma* | 0.43 | *Pcdhga8* | 5.41 | *Ccl19-ps1* |
| -0.81 | *Npy* | -0.31 | *Hnrnpc* | 0.44 | *Ppp1r1b* | 5.59 | *4930431P22Rik* |
| -0.77 | *Jun* | -0.31 | *Macroh2a1* | 0.45 | *Igfbp4* | 5.99 | *LOC100041593* |
| -0.76 | *Car12* | -0.29 | *Egr3* | 0.45 | *Zfhx4* | 6.09 | *Amd2* |
| -0.76 | *Shisa9* | -0.28 | *Stip1* | 0.45 | *Dyrk3* | 6.71 | *Ccl21c* |
| -0.73 | *Slc25a37* | -0.28 | *Khdrbs3* | 0.46 | *Kcnj4* | 6.71 | *Gm13304* |
| -0.73 | *Snurf* | -0.28 | *Calr* | 0.47 | *Tmod1* | 6.71 | *Gm10591* |
| -0.73 | *Crhbp* | -0.26 | *Hnrnph1* | 0.48 | *Ephx1* | 6.71 | *Ccl21d* |
| -0.69 | *Prokr2* | 0.27 | *Rhob* | 0.48 | *Mpp3* | 6.81 | *Gm13302* |
| -0.68 | *Pcdh8* | 0.27 | *Snn* | 0.48 | *Lurap1l* | 9.69 | *LOC100041599* |
| -0.68 | *Plscr4* | 0.27 | *Ngef* | 0.48 | *Per1* | 11.74 | *Ccl21b* |
| -0.64 | *Prss23* | 0.28 | *Camk1d* | 0.48 | *Prelid3a* | | |
| -0.64 | *Amd1* | 0.28 | *Adam23* | 0.50 | *Ngdn* | | |
| -0.63 | *Nr4a2* | 0.29 | *Slc30a3* | 0.55 | *Nptx2* | | |
| -0.63 | *Marcksl1* | 0.30 | *Fam131a* | 0.59 | *Mansc1* | | |
| -0.62 | *Pop1* | 0.30 | *Arhgef3* | 0.59 | *Armh4* | | |
| -0.61 | *Klf10* | 0.31 | *Gpr22* | 0.59 | *Camk1g* | | |
| -0.60 | *Dusp14* | 0.31 | *Diras2* | 0.60 | *Gorab* | | |
| -0.59 | *Lgi3* | 0.32 | *Tbc1d1* | 0.62 | *Lingo3* | | |
| -0.59 | *Pcdh7* | 0.32 | *Nrgn* | 0.63 | *Cpne5* | | |
| -0.57 | *Fndc3a* | 0.33 | *Hacd2* | 0.63 | *Shb* | | |
| -0.56 | *Mpdz* | 0.33 | *Sncb* | 0.65 | *Lsm11* | | |
| -0.54 | *Rnd3* | 0.34 | *Nr1d2* | 0.66 | *Hs3st1* | | |
| -0.52 | *Glt8d2* | 0.34 | *Slc7a8* | 0.68 | *Colgalt2* | | |

**All DEGs: log2fold >0.3, $p_{adj}$<0.05**

**F** DEGs *Neil1-/-Neil2-/-*

| | | | | | |
|---|---|---|---|---|---|
| -6.21 | *Gm21992* | -0.44 | *Gm40363* | 0.67 | *Camk1g* |
| -3.13 | *Fmod* | -0.44 | *Zfp950* | 0.68 | *Fstl4* |
| -2.18 | *Nlrx1* | -0.43 | *Nr4a3* | 0.70 | *Cpne5* |
| -1.75 | *Clmp* | -0.38 | *Lman2l* | 0.71 | *Pcdha12* |
| -1.75 | *Wdfy1* | -0.36 | *Hspa5* | 0.75 | *Hs3st1* |
| -1.57 | *H2ac24* | -0.36 | *R3hdm1* | 0.76 | *Slc37a4* |
| -1.36 | *Npbwr1* | -0.35 | *Rnf157* | 0.78 | *Dse* |
| -1.28 | *Htr3a* | -0.35 | *Nr4a1* | 0.81 | *Lsm11* |
| -1.25 | *Cryab* | -0.34 | *Ndst4* | 0.83 | *Nat9* |
| -1.21 | *Tmem254b* | -0.32 | *Myo9a* | 0.83 | *Mc4r* |
| -1.12 | *Npas4* | -0.30 | *Tspan3* | 0.84 | *Ift46* |
| -1.12 | *Myh7* | 0.30 | *Diras2* | 0.88 | *Dcps* |
| -1.11 | *Gfra2* | 0.31 | *Elp3* | 0.88 | *Kcnj2* |
| -1.11 | *Cntnap5b* | 0.31 | *Slc3a2* | 0.88 | *Rbm3* |
| -1.06 | *Rsc1a1* | 0.32 | *Kdm4b* | 0.93 | *Scn4b* |
| -1.06 | *Nfil3* | 0.33 | *Sec14l1* | 1.00 | *Il16* |
| -0.98 | *Esd* | 0.34 | *Vstm2l* | 1.06 | *Mid1* |
| -0.96 | *Chrna5* | 0.34 | *Orai2* | 1.21 | *Nr1d1* |
| -0.91 | *Mettl23* | 0.34 | *Ip6k2* | 1.22 | *Rgcc* |
| -0.87 | *Cldn5* | 0.34 | *Cd34* | 1.30 | *Gm2026* |
| -0.83 | *Mxra7* | 0.35 | *Slc30a3* | 1.64 | *Fxyd2* |
| -0.83 | *Lgi3* | 0.36 | *Pspc1* | 1.75 | *Sag* |
| -0.83 | *Jun* | 0.37 | *Kcnj4* | 1.93 | *Ciart* |
| -0.80 | *Bmp1* | 0.37 | *Nrp1* | 2.34 | *Gm29801* |
| -0.75 | *Scn2b* | 0.37 | *Rplp1* | 4.02 | *Gm29804* |
| -0.73 | *Dnaja4* | 0.38 | *Gpr22* | 4.46 | *Myh6* |
| -0.72 | *Kcnf1* | 0.40 | *Dipk2a* | 5.77 | *4930431P22Rik* |
| -0.68 | *Nr4a2* | 0.41 | *Igfbp4* | 6.70 | *Ccl21c* |
| -0.64 | *Pts* | 0.42 | *Nr1d2* | 6.70 | *Gm13304* |
| -0.64 | *Alg9* | 0.43 | *Lurap1l* | 6.70 | *Gm10591* |
| -0.63 | *Thsd4* | 0.44 | *Cacng8* | 6.70 | *Ccl21d* |
| -0.60 | *Sc5d* | 0.48 | *Usp2* | 8.66 | *Gm2427* |
| -0.53 | *Bace1* | 0.51 | *Nkapd1* | 9.59 | *LOC100041599* |
| -0.51 | *Ppp1ccb* | 0.51 | *Fbxo10* | 10.50 | *Ccl21b* |
| -0.50 | *Ptprg* | 0.52 | *Bhlhe41* | | |
| -0.49 | *Thpo* | 0.55 | *Cirbp* | | |
| -0.49 | *Fndc3a* | 0.58 | *Gorab* | | |
| -0.49 | *Ier5* | 0.59 | *Lingo3* | | |
| -0.49 | *Pacsin2* | 0.60 | *Ppp1r1b* | | |
| -0.49 | *Akap13* | 0.61 | *Acss1* | | |
| -0.49 | *Rcbtb2* | 0.61 | *Pcdhga6* | | |
| -0.47 | *Snurf* | 0.63 | *Gsn* | | |
| -0.46 | *Rbm28* | 0.63 | *Slc7a5* | | |
| -0.45 | *AI593442* | 0.65 | *Neo1* | | |

**All DEGs: log2fold >0.3, $p_{adj}$<0.05**

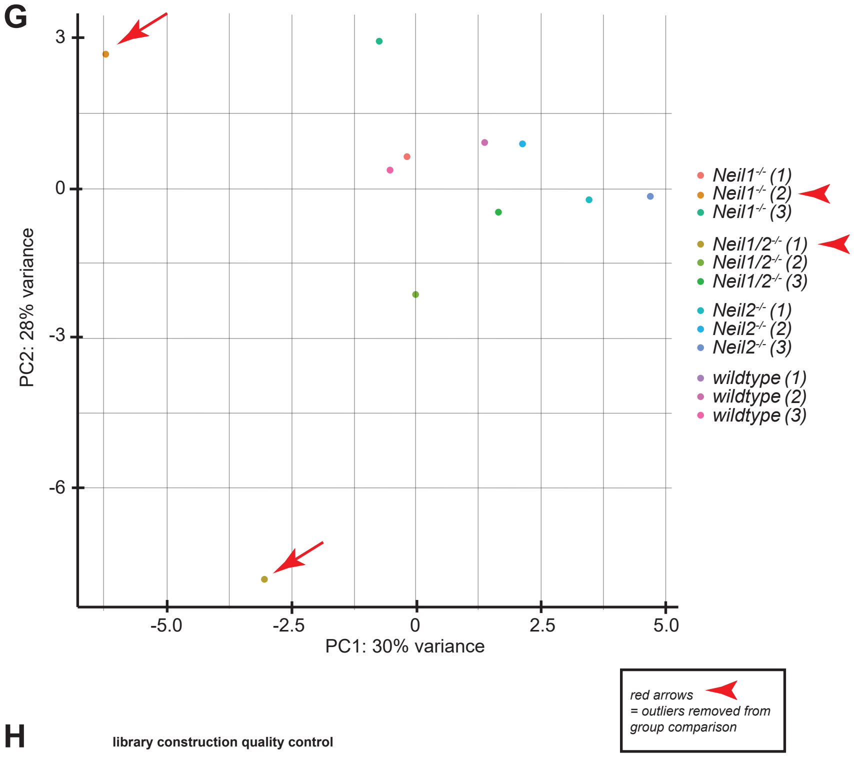


**H**

library construction quality control

| sample name | library size | mass concentration | molar concentration |
|---|---|---|---|
| wt-1 | 291 | 6.81 | 34.9 |
| wt-2 | 248 | 8.51 | 47.88 |
| wt-3 | 260 | 6.67 | 36.69 |
| N1-1 | 262 | 6.71 | 36.7 |
| N1-2 | 254 | 9.34 | 52.22 |
| N1-3 | 266 | 5.98 | 32.4 |
| N2-1 | 286 | 11.72 | 60.1 |
| N2-2 | 273 | 8.69 | 46.52 |
| N2-3 | 273 | 4.05 | 21.41 |
| N1/2-1 | 257 | 4.95 | 27.36 |
| N1/2-2 | 253 | 6.43 | 36.37 |
| N1/2-3 | 268 | 4 | 21.54 |

**Figure S4. Gene ontology biological processes enrichment analysis yielded significantly enriched terms in processes relevant for CNS function.** (A-C, colored GO-terms) GO-terms relevant for CNS function in all three genotypes (PANTHER Classification System release 15.0, analysis 8/2020, FDR<0.05, fold enrichment >2). (D-F) All DEGs for all genotypes (log2fold>0.3, padj<0.05). (G) Exploratory data analysis revealed two outliers, which were subsequently excluded from the group comparisons (red arrows). (H) Library construction quality control, outliers marked with red arrows.

#### Figure S5

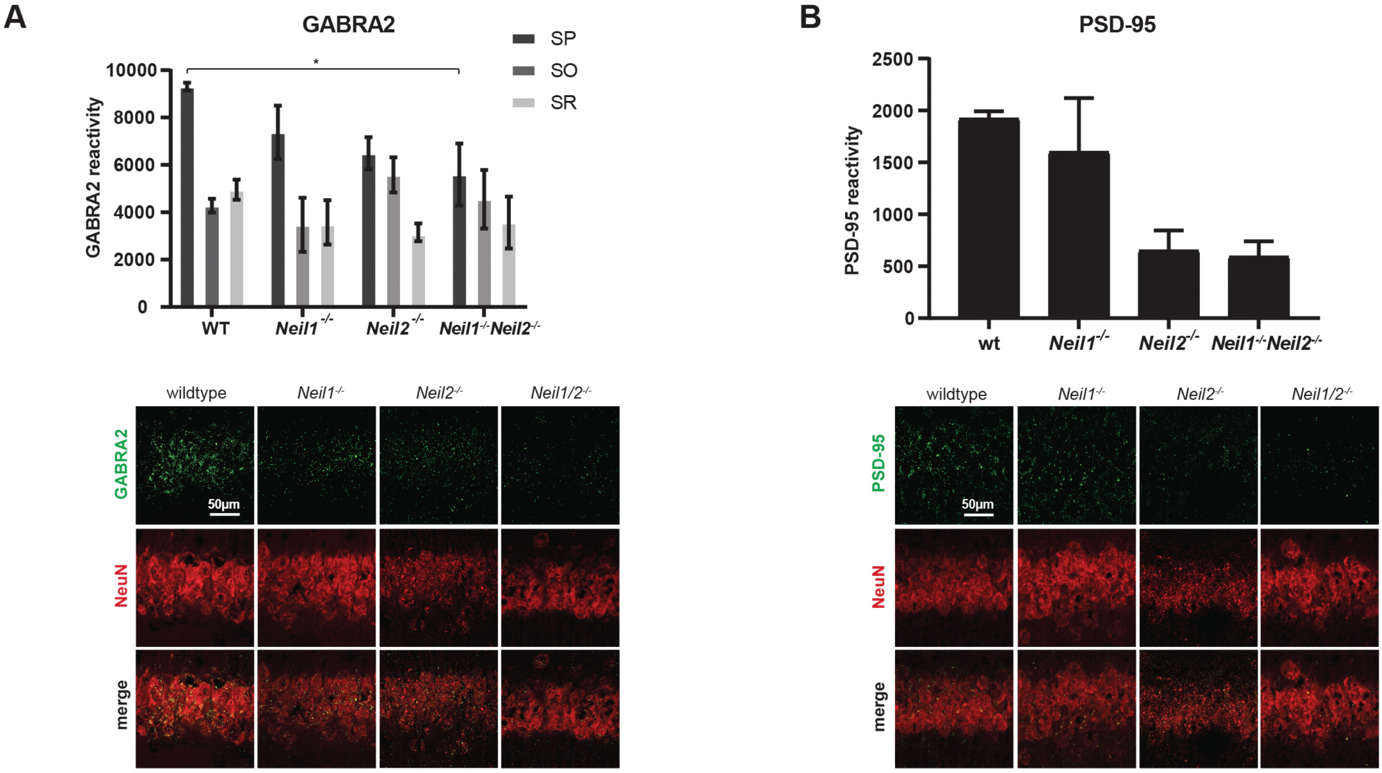


**Figure S5. Reduced expression of GABRA and PSD-95 in CA1.** (A) GABA-A-receptor α2 subunit (GABRA2) expression across CA1 stratum pyramidale (SP), stratum oriens (SO) and stratum radiatum (SR). * $p = 0.0314$ by two-way ANOVA/Tukey ($p = 0.1369$ for *Neil2*$^{-/-}$ in SP). (B) Expression of PSD-95 across SP (SR and SO were not examined due to bad immunostaining signal; $p = 0.0762$ and $0.0983$ for *Neil2*$^{-/-}$ and *Neil1*$^{-/-}$*Neil2*$^{-/-}$, respectively, by one-way-ANOVA/Tukey). (A and B) Data are presented as “reactivity levels” based on Imaris spot detection tool ± SEM. n = 6 animals (2 slices each), statistics at animal level (see methods).
